# Nutrient fluxes, oxygen consumption and fatty acid composition from deep-water demo- and hexactinellid sponges from New Zealand

**DOI:** 10.1101/2024.04.24.590875

**Authors:** Tanja Stratmann, Kathrin Busch, Anna de Kluijver, Michelle Kelly, Sadie Mills, Sven Rossel, Peter J. Schupp

## Abstract

Sponges are an important component of deep-water ecosystems enhancing eukaryotic biodiversity by hosting diverse endo- and epibiota and providing three dimensional habitats for benthic invertebrates and fishes. As holobionts they are important hosts of microorganisms which are involved in carbon and nitrogen cycling. While increasing exploration of deep-water habitats results in new sponge species being discovered, little is known about their physiology and role in nutrient fluxes. Around New Zealand (Southwest Pacific), the sponge biodiversity is particularly high, and we selected six deep-sea sponge clusters (*Saccocalyx*, *Suberites*, *Tedania*, *Halichondria*/ *Dendoricella*, Sceptrulophora, *Lissodendoryx*) for *in-situ* and *ex-situ* experiments. We investigated the biochemical composition of the sponges, measured oxygen consumption and inorganic nutrient fluxes, as well as bacterial and phospholipid-derived fatty acid (PLFA) compositions. Our aim was to assess differences in fluxes and fatty acid composition among sponge clusters and linking their bacterial communities to nitrogen cycling processes.

All sponges excreted nitrite and ammonia. Nitrate and phosphate excretion were independent of phylum affiliation (Demospongiae, Hexactinellida). Nitrate was excreted by the *Halichondria*/ *Dendoricella* and *Lissodendoryx* clusters, whereas the *Suberites*, *Tedania*, and Sceptrulophora clusters consumed it. Phosphate was excreted by Sceptrulophora and *Halichondria*/ *Dendoricella* clusters and consumed by all other clusters. Silicon was consumed by all sponge clusters, except for *Saccocalyx* and *Halichondria*/ *Dendoricella* clusters. Oxygen consumption rates ranged from 0.17 to 3.56±0.60 mmol O_2_ g C d^−1^.

The PLFA composition was very sponge-cluster dependent and consisted mostly of long-chain fatty acids. Most PLFAs were sponge-specific, followed by bacteria-specific PLFAs, and others. All sponge clusters, except for *Suberites*, were low-microbial abundance (LMA) sponges whose bacterial community composition was dominated by Proteobacteria, Bacteroidota, Planctomycetota, and Nitrospinota. The *Suberites* cluster consisted of high-microbial abundance (HMA) sponges with Proteobacteria, Chloroflexota, Acidobacteriota, and Actinobacteriota as dominant bacteria.

Based on the inorganic nitrogen flux measurements, we identified three types of nitrogen cycling in the sponges: In type 1, sponges (*Dendoricella* spp. indet., *Lissodendoryx* cluster) respired aerobically and ammonificated organic matter (OM) to ammonium, fixed N_2_ to ammonium, and nitrified aerobically heterotrophically produced ammonium to nitrate and nitrite. In type 2, sponges (*Halichondria* sp., Sceptrulophora, *Suberites*, *Tedania* clusters) respired OM aerobically and ammonificated it to ammonium. They also reduced nitrate anaerobically to ammonium via dissimilatory nitrate reduction to ammonium. In type 3, ammonium was microbially nitrified to nitrite and afterwards to nitrate presumably by ammonium-oxidizing Bacteria and/ or Archaea.

## 1. Introduction

Sponges (phylum Porifera) are the evolutionary oldest metazoans on our planet (Feuda et al., 2017; Müller et al., 2007; Simion et al., 2017) and important members of benthic ecosystems (Bell, 2008). In the Queen Charlotte Basin at the western continental shelf of Canada (NE Pacific), for instance, glass sponges (class Hexactinellida) form sponge reef complexes with a total area of 182 km^2^ and a maximum reef height of 21 m (Conway et al., 2005). Radiocarbon dating of mollusk shells indicated that these reef complexes are between 1,430±50 and 5,700±60 years old (Conway et al., 1991). They host a diverse megafauna, including crustaceans, such as crabs, shrimps, prawns, and euphausiids, and rockfish (Cornway et al., 2001). In soft-sediment abyssal plains, glass sponge stalks can be considered “habitat islands” that provide hard substrate for other deep-sea epifauna (Beaulieu, 2001). The analysis of 2,418 sponge stalks in photographs taken at Station M (NE Pacific) revealed that all epifauna were facultative suspension feeders that belonged mostly to zoanthids, tunicates, ophiuroids, and actinarians (Beaulieu, 2001). In tropical coral reefs, sponges are often the most diverse benthic phylum and in Caribbean coral reefs, they belong to the four organism groups with the largest areal coverage besides algae, scleractinian corals, and octocorals (Diaz and Rützler, 2001). In open reef habitats that are exposed to light, sponges cover up to 24% of the hard substrate (Zea, 1993), whereas in cryptic sub-rubble habitats that receive less light, they cover up to 54% of the area (Meesters et al., 1991).

Depending on their biomass, sponges contribute significantly to benthic respiration. It was estimated that the whole deep-water sponge *Geodia barretti* Bowerbank, 1858 population in the 300 km^2^ large Træna Marine Protected Area (Norwegian continental shelf) respires 60 t carbon (C) d^−1^ (Kutti et al., 2013). In fact, the entire sponge ecosystem at the Norwegian continental shelf is estimated to respire 306,000 tC yr^−1^ (Cathalot et al., 2015). Furthermore, in oligotrophic areas, sponges can also be important sources of nitrogen (N) (Bell, 2008). Sponges in Mediterranean sublittoral rocky bottom habitats were estimated to release 2.5 to 7.9 mmol N m^−1^ d^−1^ (Jiménez and Ribes, 2007), whereas at Marion Lagoon (Western Australia), Keesing et al. (2013) calculated a N release by sponges of 0.35 to 0.63 mmol N m^−1^ d^−1^. This corresponds to 10 to 18% of the recycled N in the oligotrophic west Australian continental shelf (Keesing et al., 2013). Sponges are also a major transformer of dissolved organic matter (DOM) (Hildebrand et al., 2022).

Sponges are holobionts (Webster and Taylor, 2012) consisting of the sponge itself and a microbial community that can contribute up to 38% to the overall sponge holobiont volume (Vacelet, 1975). Based on the number of microorganisms present in the sponge holobiont, they are classified as high-microbial-abundance (HMA) sponges or low microbial abundance (LMA) sponges. In shallow waters, low-microbial-abundance sponges contain typically around 10^5^ to 10^6^ bacteria g^−1^ sponge tissue which is comparable to the density of microorganisms in seawater, whereas HMA sponges typically host around 10^8^ to 10^10^ microorganisms g^−1^ sponge tissue (Hentschel et al., 2006). Shallow-water LMA sponges often have one dominating bacterial phylum (Giles et al., 2013), whereas shallow-water HMA sponges are dominated by a more diverse bacterial community (Erwin et al., 2015; Moitinho-Silva et al., 2017). Among others, they are Chloroflexota hotspots (Bayer et al., 2018). In the deep sea, LMA glass sponges (Hexactinellida) host a microbiome that is different from the microbiome of LMA demosponges, HMA sponges or reference water samples (Busch et al., 2022; Steinert et al., 2020).

Fatty acids are components of lipids and serve as source of energy (Lindsay, 1975) and building blocks of membranes (Spector and Yorek, 1985; van Deenen, 1966). Furthermore, they are important for transducing signals in cells (Faergeman and Knudsen, 1997; Graber et al., 1994) and for the regulation of gene expression (Sampath and Ntambi, 2004). As several fatty acids can only be synthesized by specific taxa, these may be used as dietary tracers and provide information about diet preferences and food origin (Kelly and Scheibling, 2012). Bacteria, for instance, produce fatty acids with iso (*i*) and anteiso (*ai*)-branched carbon (C) chains, such as *i*-C15:0, *ai*-C15:0, *i*-C17:0, and *ai*-C17:0 (Kaneda, 1991). Sponges, in comparison, produce long chain fatty acids (≥C24) via elongation of shorter chain fatty acids they take up as part of their diet (Carballeira et al., 1986; de Kluijver et al., 2021). These fatty acids are known as “demospongic acids” because they were first discovered in demosponges (Litchfield et al., 1976), though they are also present in glass sponges, but not in calcareous sponges (Lawson et al., 1984; Schreiber et al., 2006). As sponge holobionts consist of both, sponge cells and microorganisms, the fatty acid composition of sponges usually includes bacteria- and sponge-specific fatty acids (e.g., Bart et al., 2020; de Kluijver et al., 2021; Morganti et al., 2022; Rix et al., 2016; van Duyl et al., 2020).

In this study, we measured *in-situ* and *ex-situ* oxygen and inorganic nutrient fluxes in deep-sea sponges (595 to 4161 m water depth) around New Zealand in the Southeast Pacific. New Zealand belongs to the “temperate Australasia” biogeographic realm (Spalding et al., 2007) which has one of the highest sponge biodiversities (1361 sponge species; (Kelly and Sim-Smith, 2023; Spalding et al., 2007; Van Soest et al., 2012)). We investigated whether oxygen and nutrient fluxes differed between sponge clusters (i.e., sponges for which a species level distinction was not possible based on DNA barcoding and MALDI-TOF MS), and we tried to link bacterial communities in sponges with inorganic nutrient fluxes to decipher dominant nitrogen cycling processes inside the sponges. Furthermore, we assessed differences in the fatty acid composition among the various sponge clusters.

## 2. Materials and methods

### 2.1 *In-situ* incubation experiment

During R/V *Sonne* cruise SO254 around New Zealand (Fig. 1) in January and February 2017, a sponge (TS001, Fig. 2, Table 1, Fig. S1) was incubated *in-situ* at 4161 m in a 50×50×50 cm large benthic incubation chamber (CUBE; (Stratmann et al., 2018)) for 5 h. For this purpose, the CUBE was placed over the sponge by the remotely operated vehicle ROV Kiel 6000 (GEOMAR, Kiel, Germany) and water samples (∼35 ml) were taken automatically every hour for the measurement of inorganic nutrient concentrations (nitrate, nitrite, ammonium, phosphate, silicon). Oxygen concentration (µmol O_2_ kg^−1^ seawater) was recorded continuously every 10 sec by an oxygen optode (Contros HydroFlash® O_2_; Kongsberg Maritime Contros GmbH, Germany) which had been calibrated by 2-point calibration (0% calibration: Na_2_SO_3_; 100% calibration: air-bubbled seawater). A second CUBE was placed over bare sediment in close proximity and served as control/ blank. This CUBE took water samples automatically at the same time intervals as the sponge incubation CUBE and measured oxygen concentration continuously. At the end of the incubation, the sponge was collected by the ROV and returned to the vessel in a closed biobox. Aboard, the length *L*, width *W*, and height *H* of the sponge was measured with a ruler to the nearest 0.1 cm and a subsample of sponge tissue was taken with a scalpel, rinsed in sterile seawater, flash-frozen in liquid nitrogen and stored at −80°C for microbial analysis. An additional tissue sample was taken for species identification before the remaining sponge was frozen at - 21°C. Water samples for nutrient analysis were filtered through 0.45 µm filters into 10 ml vials and stored frozen (−21°C).

**Figure 1.**
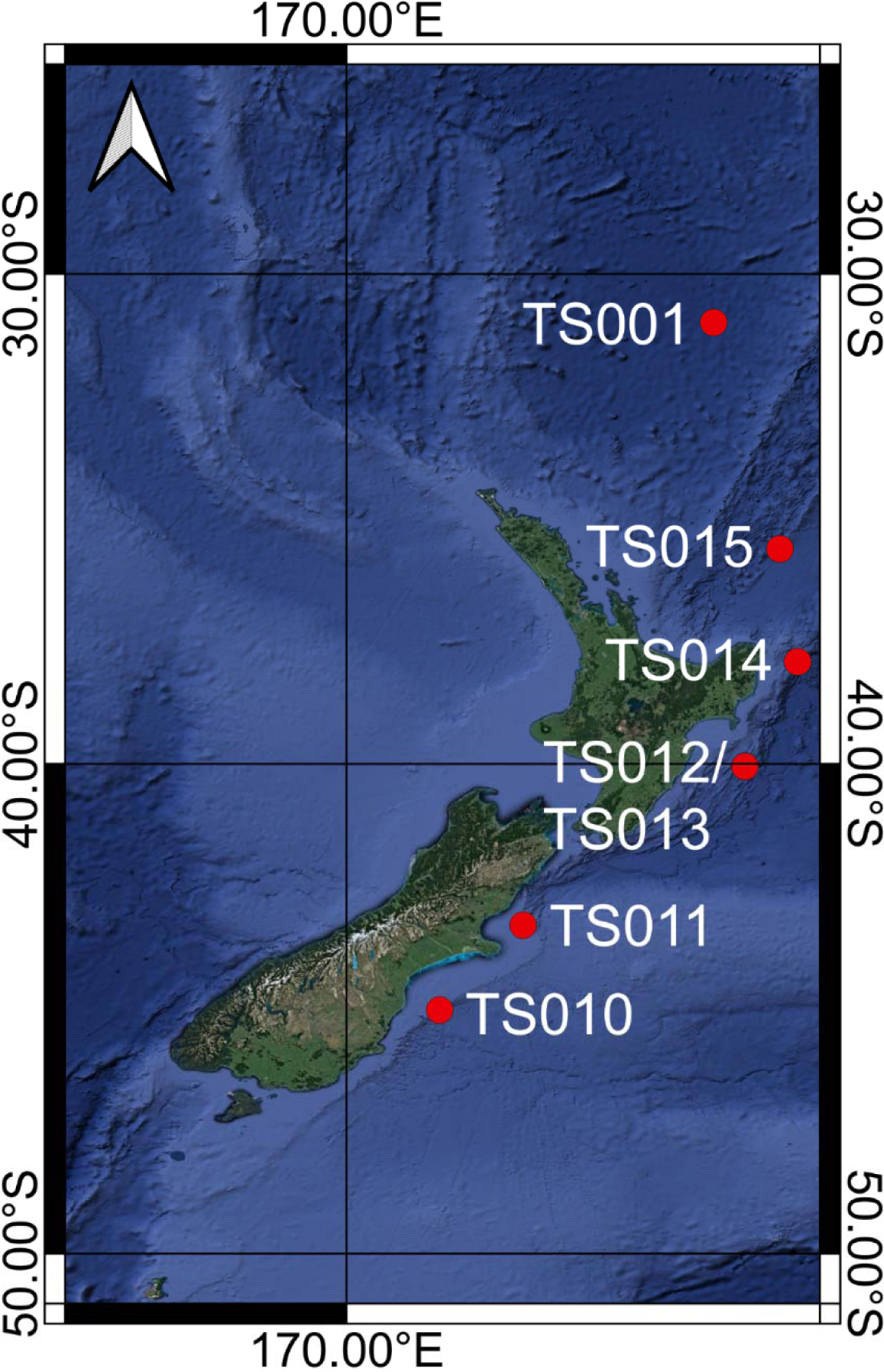
Map with sampling sites (red dots) off New Zealand where the deep-sea sponges were collected for the incubations. Detailed information about species names and sampling depths are given in Table 1.

**Figure 2.**
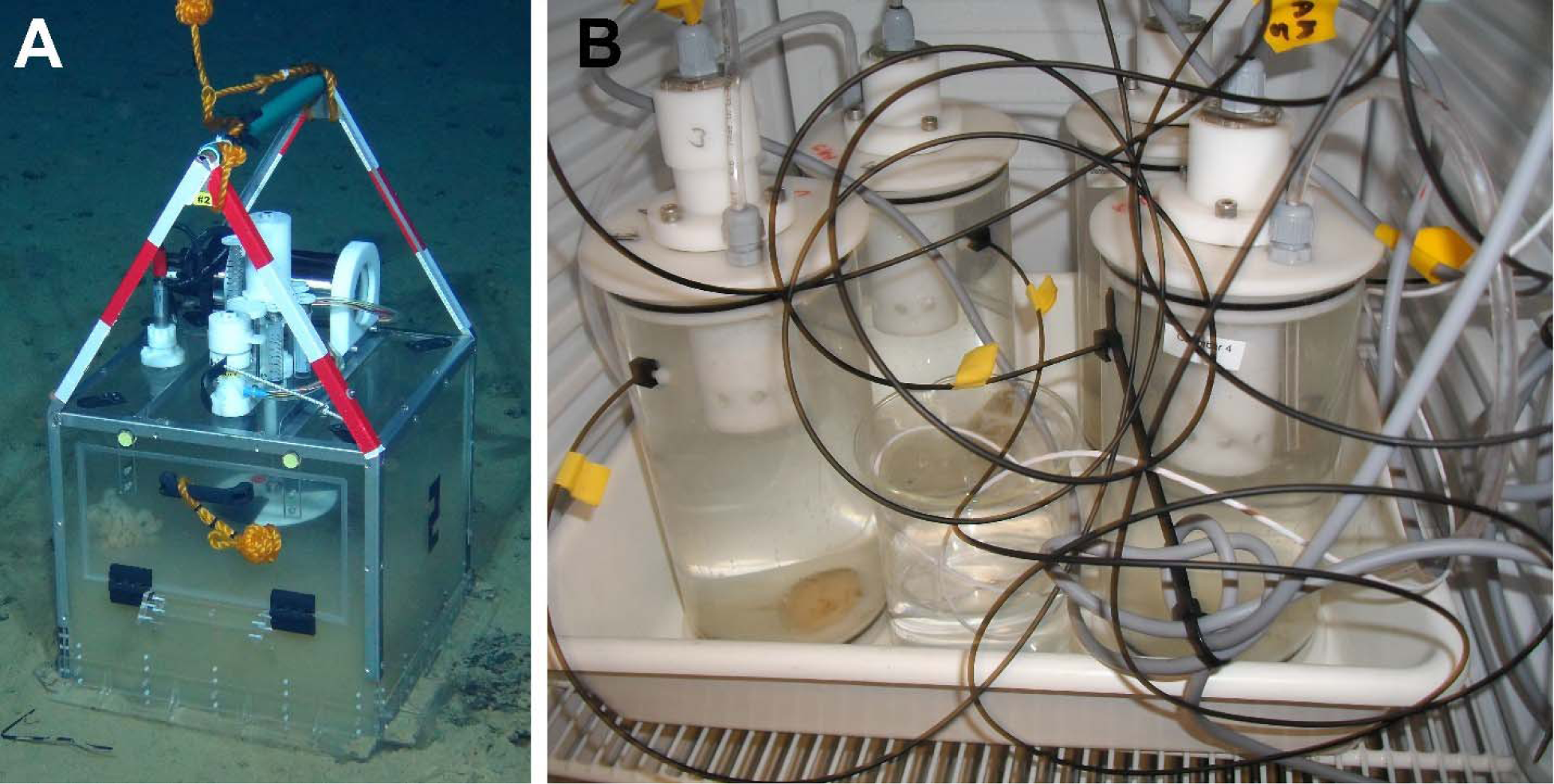
Incubation of deep-sea sponges (A) *in-situ* at ∼4,000 m water depth and (B) *ex-situ* aboard R/V Sonne. *In-situ* photo was taken with ROV Kiel 6000 (GEOMAR, Kiel, Germany) and *ex-situ* photo was taken by Tanja Stratmann.

**Table 1.**
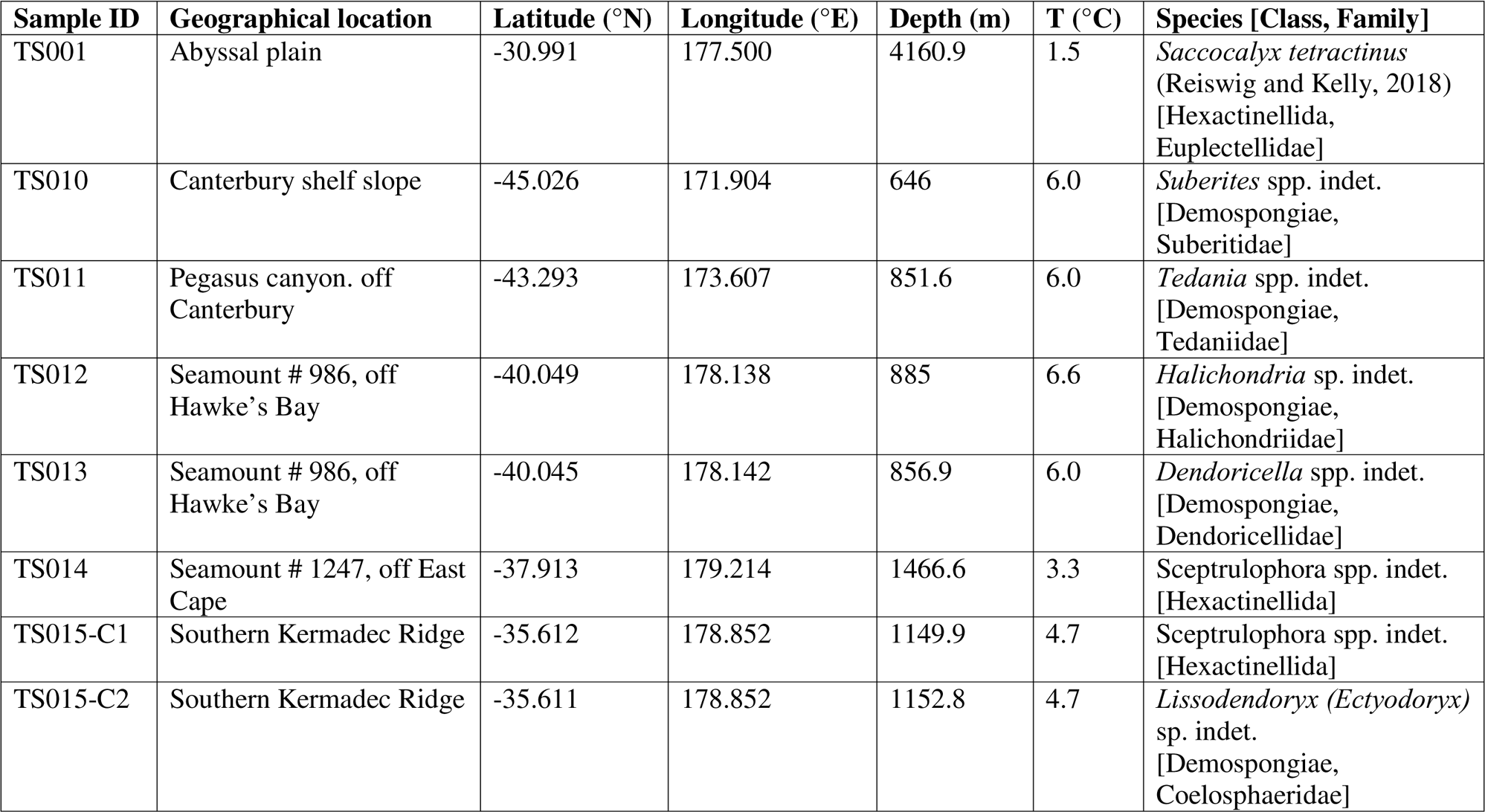
Detailed information about sampling location and species name of all collected sponges. Abbreviation: T = water temperature.

| Sample ID | Geographical location | Latitude (°N) | Longitude (°E) | Depth (m) | T (°C) | Species [Class, Family] |
| --- | --- | --- | --- | --- | --- | --- |
| TS001 | Abyssal plain | -30.991 | 177.500 | 4160.9 | 1.5 | *Saccocalyx tetractinus*<br>(Reiswig and Kelly, 2018)<br>[Hexactinellida,<br>Euplectellidae] |
| TS010 | Canterbury shelf slope | -45.026 | 171.904 | 646 | 6.0 | *Suberites* spp. indet.<br>[Demospongiae,<br>Suberitidae] |
| TS011 | Pegasus canyon. off<br>Canterbury | -43.293 | 173.607 | 851.6 | 6.0 | *Tedania* spp. indet.<br>[Demospongiae,<br>Tedaniidae] |
| TS012 | Seamount # 986, off<br>Hawke's Bay | -40.049 | 178.138 | 885 | 6.6 | *Halichondria* sp. indet.<br>[Demospongiae,<br>Halichondriidae] |
| TS013 | Seamount # 986, off<br>Hawke's Bay | -40.045 | 178.142 | 856.9 | 6.0 | *Dendoricella* spp. indet.<br>[Demospongiae,<br>Dendoricellidae] |
| TS014 | Seamount # 1247, off East<br>Cape | -37.913 | 179.214 | 1466.6 | 3.3 | *Sceptrulophora* spp. indet.<br>[Hexactinellida] |
| TS015-C1 | Southern Kermadec Ridge | -35.612 | 178.852 | 1149.9 | 4.7 | *Sceptrulophora* spp. indet.<br>[Hexactinellida] |
| TS015-C2 | Southern Kermadec Ridge | -35.611 | 178.852 | 1152.8 | 4.7 | *Lissodendoryx (Ectyodoryx)*<br>sp. indet.<br>[Demospongiae,<br>Coelosphaeridae] |

### 2.2 *Ex-situ* incubation experiment

During R/V *Sonne* cruise SO254 sponges were collected at water depths between 595 m and 1,467 m (Table 1) with ROV Kiel 6000. Aboard, individual sponges of the same putative species were incubated for 5 h in three closed incubation chambers (cylinder height: 17 cm, diameter: 10 cm, volume: 1.26 L; Fig. 2) at *in-situ* temperature in the dark. A fourth incubation chamber filled with ambient seawater served as control/ blank. Before the start (T0) and at the end of the incubations (T5), duplicate water samples were taken from each incubation chamber for inorganic nutrient concentration, filtered (0.45 µm filters), and stored frozen (−21°C). During the incubations, seawater air saturation (% O_2_) was recorded continuously in 1 s intervals using an optical oxygen meter (FireStingO_2_, PyroScience GmbH, Germany) after 2-point optode calibration. After completing the incubations, length *L*, width *W*, and height *H* of each sponge specimen were measured and tissue subsamples for microbial community analysis were taken from several specimens (Table S1) with a scalpel. These samples were rinsed in sterile seawater, flash-frozen in liquid nitrogen, and stored at −80 °C. The remaining sponge tissue was frozen at - 21°C after another tissue subsample was taken from one specimen per putative species for species identification.

### 2.3 Sample processing

#### 2.3.1 Species identification

One sponge specimen per putative species was identified by spicule analysis or, in case of *Lissodendoryx (Ectyodoryx)* sp. indet., based on still photo analysis. Voucher specimens are deposited in the NIWA Invertebrate Collection, Wellington, New Zealand. Additionally, small samples of sponge tissue were taken from each sponge specimen for DNA barcoding and matrix assisted laser desorption/ionisation - time of flight mass spectrometry (MALDI-TOF MS).

For DNA-barcoding, sponge tissue was incubated in 30 µl Chelex (InstaGene^TM^ Matrix, BioRad, USA) for 50 min at 56°C and 10 min at 96°C. In a total reaction volume of 20µl, 2 µl DNA extract was mixed with 0.2µl of forward (*16S1fw*) and reverse (*16SH_mod*) primers (Dohrmann et al., 2008), 10 µl Accu Start PCR mix (2xPCR master mix, Quantabio, USA), and 7.6 µl molecular grade water. The PCR protocol was adapted from Kersken et al., (2018) using the following settings: initial denaturation for 5 min at 94°C followed by 40 cycles of denaturation at 94°C for 30 s, annealing for another 30 s at 48°C, and elongation at 72°C for 45 s. This was followed by a final elongation for 3 min at 72°C. Sanger sequencing was carried out at Macrogen Europe (Amsterdam, Netherlands), producing chromatograms for forward and reverse sequences for each specimen. These chromatograms were processed in the *Geneious R7* software (version 7.0.6; https://www.geneious.com) to produce consensus sequences which were blasted (Altschul et al., 1997) to avoid contaminations. Final sequences were aligned in the *SeaView* software (Gouy et al., 2010) using the muscle algorithm and by-eye control. At the end, a neighbor joining analysis was carried out using Kimura two-parameter (K2P) distances to check for sequence similarities.

For MALDI-TOF MS, sponge tissue was incubated in a saturated solution of α-Cyano4-hydroxycinnamic acid (HCCA) dissolved in 50% acetonitrile, 47.5% molecular grade water, and 2.5% trifluoroacetic acid that completely covered the tissue as suggested by Rossel et al., (2024). After the incubation, 1.5 µl extract was transferred to a target plate and measured using a Microflex LT/SH System (Bruker Daltonics, USA). Molecule masses were measured from 2 to 20k Dalton (kDa) applying the *flexControl 3.4.* software (Bruker Daltonics). Peaks of a mass range between 2 and 20 kDa and a peak resolution of >400 were evaluated using the centroid peak detection algorithm with a signal-to-noise threshold of two and a minimum intensity threshold of 600. The proteins/oligonucleotide method was employed at a maximal resolution of 10 times above the threshold to validate fuzzy control. To create a sum spectrum, a total of 120 laser shots were applied to a spot which was measured three times each. Resulting mass spectra were processed with the *MALDIquantForeign* package (version 0.13) (Gibb, 2015) and the *MALDIquant* package (version 1.22) (Gibb and Strimmer, 2012) in *R* (version 4.2.2) (R-Core Team, 2022). Spectra were square-root transformed, smoothed using the Savitzky Golay method (Savitzky and Golay, 1964), baseline corrected using the SNIP method (Ryan et al., 1988), and spectra normalized using the total ion current (TIC) method. Repeated measurements were averaged using mean intensities. Peak picking was carried out using a signal to noise ratio (SNR) of 5 and a half window size of 17. Repeated peaks were binned to align homologous mass peaks. The resulting data were Hellinger transformed (Legendre and Gallagher, 2001) and hierarchically clustered in *R* using Ward’s *D* and Euclidean distances.

Based on the DNA-barcoding and MALDI-TOF MS results (Supplementary Material; Fig. S2), sponges were classified in six sponge clusters (Fig. 3).

**Figure 3.**
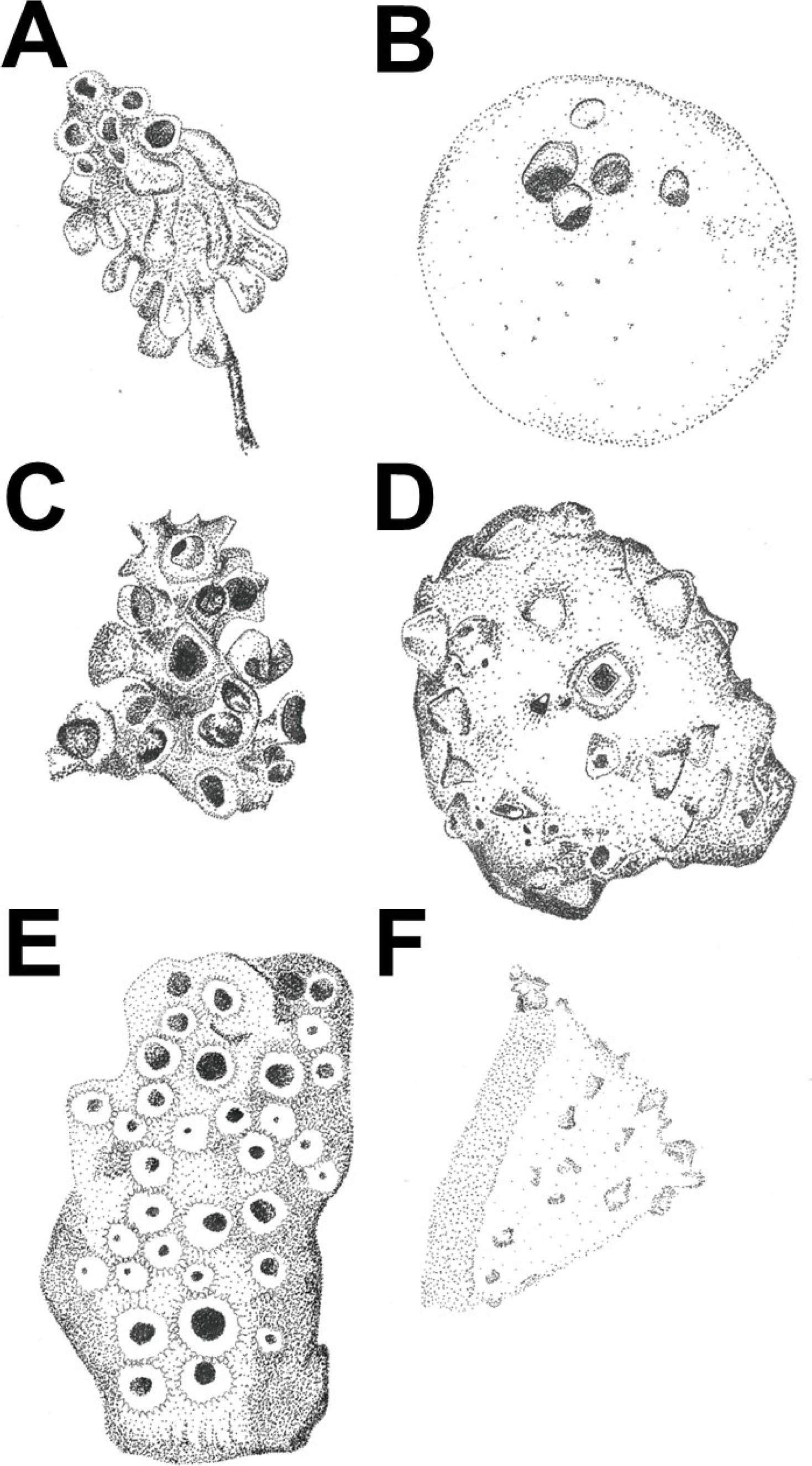
Drawings of the six sponge clusters A) *Saccocalyx* cluster, B) *Suberites* cluster, C) Sceptrulophora cluster, D) *Tedania* cluster, E) *Halichondria*/ *Dendoricella* cluster, F) *Lissodendoryx* cluster. Sizes of sponge clusters in the drawing are not to scale. Illustrations by Tanja Stratmann.

#### 2.3.2 Sponge tissue analysis

Sponge volume (*V*, cm^3^) was determined via water displacement (*V_WD_*) and based on its geometric shape (*V_GS_*). For the latter, each sponge was classified into different three-dimensional geometric shapes (cuboid, sphere, cylinder, circular cone, frustum; Fig. S3) and *V_GS_* was calculated as:

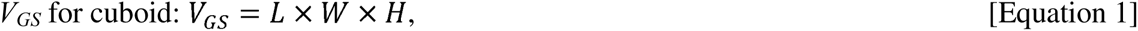

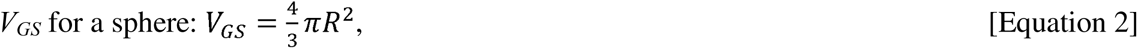

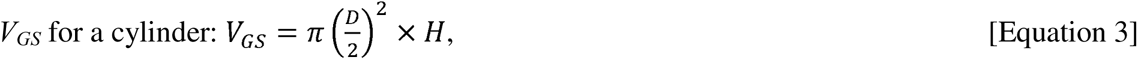

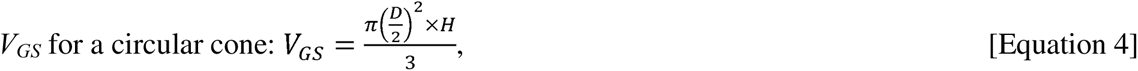

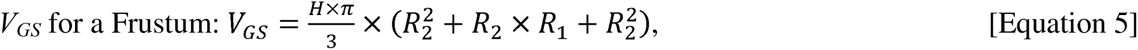

where *L* = sponge length, *W* = sponge width, *H* = sponge height, *R* = sponge radius, *D* = diameter of the sponge, and *R_1_* and *R_2_* correspond to the two different radii in Fig. S3. The *V_GS_*:*V_WD_*-ratio for the different sponges species clusters is presented in Table S3.

Sponge wet mass (*WM*, g) of remaining tissue was measured with a balance to the nearest 0.001 g. Dry mass (*DM*, g) was determined after freeze-drying and ash-free dry mass (*AFDM*, g) was measured after oxidizing sub-samples (0.1 g) of freeze-dried sponges in a muffle furnace at 450°C for 4 h. Total carbon (*TC*, % DM)/ δ^13^TC content and total nitrogen (*TN*, % DM)/ δ^15^N content of ∼20 mg freeze-dried, finely-ground, homogenized sponge tissue samples were measured with a Thermo Flash EA 1112 elemental analyzer (EA; Thermo Fisher Scientific, USA) coupled to a DELTA V Advantage Isotope Ratio Mass Spectrometer (IRMS; Thermo Fisher Scientific, USA). To study organic carbon (org. C, % DM)/ δ^13^org. C content of the sponges, ∼20 mg tissue powder was packed in 8×5 mm pre-combusted (4 h at 450°C) silver capsules, acidified with 20 µL 2% HCl, and dried at 60°C on a hot plate. Afterwards, capsules were closed and measured with the same EA-c-IRMS.

Total *WM* (g) of each incubated sponge specimen was determined based on *V_GS_* using the onboard size measurements of the sponges and conversion factors from Table S3, because no sponge arrived intact in the Netherlands due to the subsampling aboard.

Sponge condition index *CI* (−) was calculated following Lüskow et al., (2019):

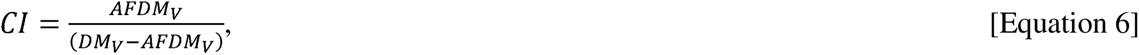

where *AFDM_V_* is the volume specific ash-free dry mass (g ADFM cm^−3^ sponge) and *DM_V_* is the volume specific dry mass (g DM cm^−3^ sponge).

#### 2.3.3 Fatty acid extraction and analysis

PLFAs were extracted from ∼100 mg freeze-dried, finely-ground, homogenized sponge tissue following a modified version of the Bligh and Dyer extraction method (Bligh and Dyer, 1959; de Kluijver, 2021; de Kluijver et al., 2021): Total lipids from sponge tissue were extracted in sampling tubes with 15 ml methanol (MeOH), 7.5 ml dichloromethane (DCM), and 6 ml phosphate (P)-buffer (8.7 g K_2_HPO_4_ dissolved in 1 l MilliQ-water, pH adjusted to 7–8 with 1 mol l^−1^ HCl) during constant shaking for 3 h. After addition of 7.5 ml DCM and 7.5 ml P-buffer, layers separated by leaving the sample tubes in the freezer (−21°C) over night. The DCM layer was transferred to new tubes and 7.5 ml DCM were again added to the original tube. After the layers separated via centrifugation with 170 *g* for 3 min at room temperature, the DCM layer was transferred to the same new tube and evaporated to complete dryness in a TurboVap evaporator under gentle N_2_-stream. 1 ml of DCM was added and evaporated again to complete dryness, before the sample was dissolved in 1 ml DCM:MeOH (1:1), transferred to a new vial, and dried at a FlexiVap™ Work Station under gentle N_2_ stream.

To separate the Bligh and Dyer extract into the different polarity classes, the sample was dissolved in 0.5 ml DCM and added to a column with silica acid gel (activated at 120°C for 2 h). Subsequently, the sample on the silica column was eluted with 7 ml acetone and 15 ml MeOH. The latter fraction contained the phospholipids and was therefore collected and dried in the TurboVap evaporator.

Phospholipids were derivatized to fatty acid methyl esters (FAMEs) via mild alkaline methylation. For this, the dried sample was dissolved in 1 ml MeOH/ toluene (1:1 v/v) and 1 ml 0.2 mol l^−1^ methanolic NaOH. 50 µL 0.1 mg ml^−1^ internal C_19:0_-FAME standard was added and the whole solution was incubated for 15 min at 37°C. The methylation was stopped by adding 2 ml hexane, 0.3 ml 1 mol l^−1^ acetic acid, and 2 ml Milli-Q water to the vial and shaking well. When the layers had separated, the hexane-containing upper layer was transferred to a new vial and this step was repeated twice before the hexane was evaporated until complete dryness. 50 µl 0.1 mg ml^−1^ internal C_12:0_-FAME standard was added, 100 µl hexane was added and the solution was transferred to a GC analysis vial.

All samples were stored at −20°C until FAMEs concentrations were measured on a gas chromatograph (GC) with flame ionization detector (FID) (HP 6890 series) on a non-polar analytical column (Agilent, CP-Sil5 CB; 25 m x 0.32 mm x 0.12 µm). The retention times of the individual peaks on the gas chromatogram were converted to equivalent chain length (ECL) using the retention times of the standard C_12:0_-FAME, C_16:0_-FAME, and C_19:0_-FAME. The concentration of every individual FAMEs (µg g dry mass^−1^) was quantified using the known concentration of C_12:0_-FAME.

#### 2.3.4 Analysis of bacterial communities of sponges

Ten sponge specimens were analyzed for their bacterial community composition (Table S1). DNA was extracted of ∼0.25 g of sponge tissue and its quality and quantity were checked by NanoDrop and gel electrophoresis after a PCR with universal 16S rRNA gene primers. Then, the bacterial V3 to V4 variable regions were amplified in a one-step PCR, using the primer pair 341F-806R (Caporaso et al., 2011; Muyzer et al., 1993). Before sequencing the bacterial libraries on a MiSeq platform (MiSeqFGx, Illumina) with v3 chemistry, a quality check was performed by gel electrophoresis, normalization, and pooling.

Amplicon sequences were processed within QIIME2 (version 2019-10) (Bolyen et al., 2019). Demultiplexed forward reads were imported into QIIME2 and primers were trimmed. Amplicon sequence variants (ASVs) were generated from single-end reads (truncated to 270 nt) with the DADA2 algorithm (Callahan et al., 2016). Representative ASVs were taxonomically classified with the SILVA database (version 138 99% OTUs 16S) (Quast et al., 2013), using bacterial primer specific trained Naïve Bayes taxonomic classifiers.

#### 2.3.5 Inorganic nutrient concentration measurements and flux calculations

Water samples for inorganic nutrient analysis were thawed for 24 h prior to the measurements of ammonium, nitrite, nitrate, silicon, and phosphate with a SEAL QuAAtro analyzer (Bran+Luebbe, Germany).

For the *ex-situ* incubations, blank- and biomass-corrected fluxes (*F_i.nut_*) of the inorganic nutrients (µmol g C^−1^ d^−1^) were calculated as follows:

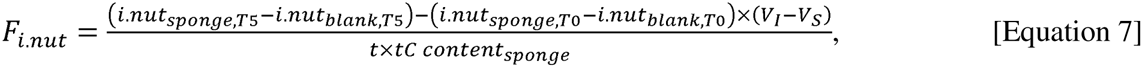

where *i.nut_sponge_* is the inorganic nutrient concentration (µmol l^−1^) in the incubation chamber with the sponge. *i.nut_blank_* is the inorganic nutrients concentration in the incubation chamber filled with ambient seawater (*blank*). *T0* corresponds to the start of the incubation and *T5* corresponds to the end of the incubation after 5 h. *t* is the incubation time (=0.21 d) and *tC content_sponge_* is the total org. C content of the incubated sponge (g C). *V_I_* is the volume of the incubation chamber (= 1.26 l) and *V_S_* is the volume of the sponge (l).

For the *in-situ* incubation, fluxes (*F_i.nut_*) of the inorganic nutrients (µmol g C^−1^ d^−1^) were calculated as follows:

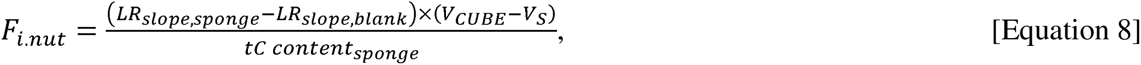

where *LR_slope,sponge_* and *LR_slope,water_* are the estimated slopes of the linear regression analyses performed with the concentrations of inorganic nutrients (µmol l^−1^) collected over 5 hours during the *in-situ* incubations as response (dependent) variable and time as predictor (independent) variable. *tC content_sponge_* is the total org. C content of the incubated sponge (g C). *V_CUBE_* is the volume of the CUBE (= 125 l) and *V_S_* is the volume of the sponge (l).

A list with inorganic nutrient concentrations at the begin of the *in-situ* and *ex-situ* incubation experiments is shown in Table S2.

#### 2.3.6 Oxygen consumption calculations

Air saturations recorded by the oxygen meter in the *ex-situ* experiments were converted to absolute oxygen concentrations (µmol O_2_ l^−1^) following (Weiss, 1970) using the *marelac* package (Soetaert et al., 2010) in *R* (R-Core Team, 2022). Subsequently, average decrease in oxygen over time (ΔO_2_, mmol O_2_ l^−1^ d^−1^) in the *in-situ* and *ex-situ* experiments was calculated by linear regression and blank and biomass-corrected metabolic rate *R* (mmol O_2_ g C^−1^ d^−1^) was calculated as:

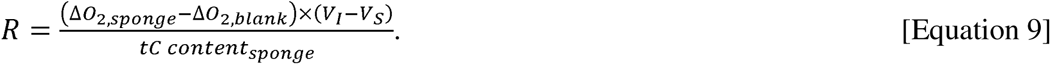

### 2.4 Statistical analysis

It was tested whether average inorganic nutrient and oxygen fluxes of the same sponge cluster differed significantly from 0 µmol g C^−1^ d^−1^ using a 1-sided Student’s t-test (α = 0.05) in *R* (version 4.3.0) after normality of data was confirmed with a Shapiro-Wilk normality test. When data were not normally distributed, a 1-sample Wilcoxon test (α = 0.05) was performed in *R*. Statistical differences in microbial community composition between the *Halichondria*/ *Dendoricella* cluster, the *Tedania* cluster, and the Sceptrulophora cluster were investigated by pairwise PERMANOVAs using the R package *vegan* (Oksanen et al., 2017). The microbial community composition of the *Lissodendoryx* cluster, the *Suberites* cluster, and the *Saccocalyx* cluster were not assessed statistically, because this parameter was studied in n < 2 specimens per cluster.

All data are presented as mean ± standard error.

## 3. Results

### 3.1 Composition of sponge tissue

Sponges consisted of 82.4 ± 3.91% water (n = 18) and their dried tissue contained 5.47 ± 0.75% TC, 4.47 ± 0.52% org. C, and 1.06 ± 0.15% TN (n = 15). The *Halichondria*/ *Dendoricella* cluster (n = 5) had the lowest TC (3.25 ± 0.35%), org. C (2.86 ± 0.39%), and TN contents (0.65 ± 0.08%), whereas the *Tedania* cluster (n = 2) had the highest TC (9.21 ± 2.19%), org. C (7.17 ± 0.31%), and TN contents (1.87 ± 0.43%). Sponge *CI* ranged from 0.11 ± 0.02 for the *Tedania* cluster (n = 3) to 0.34 ± 0.00 for the *Suberites* cluster (n = 3). The mean δ^13^org. C-value of the sponges was −17.6 ± 0.57‰ [min: −20.1 ± 0.35‰, Sceptrulophora cluster, n = 4; max: - 14.1‰, *Saccocalyx* cluster, n = 1), the mean δ^13^TC-value was −19.0 ± 0.28‰ [min: - 20.3 ± 0.09‰, Sceptrulophora cluster, n = 4; max: −17.0 ± 0.24‰, *Tedania* cluster, n = 2), and the mean δ^15^N-value was 18.5 ± 1.28‰ [min: 10.2‰, *Lissodendoryx* cluster, n = 1; max: 23.5‰, *Saccocalyx* cluster, n = 1) (Fig. 4).

**Figure 4.**
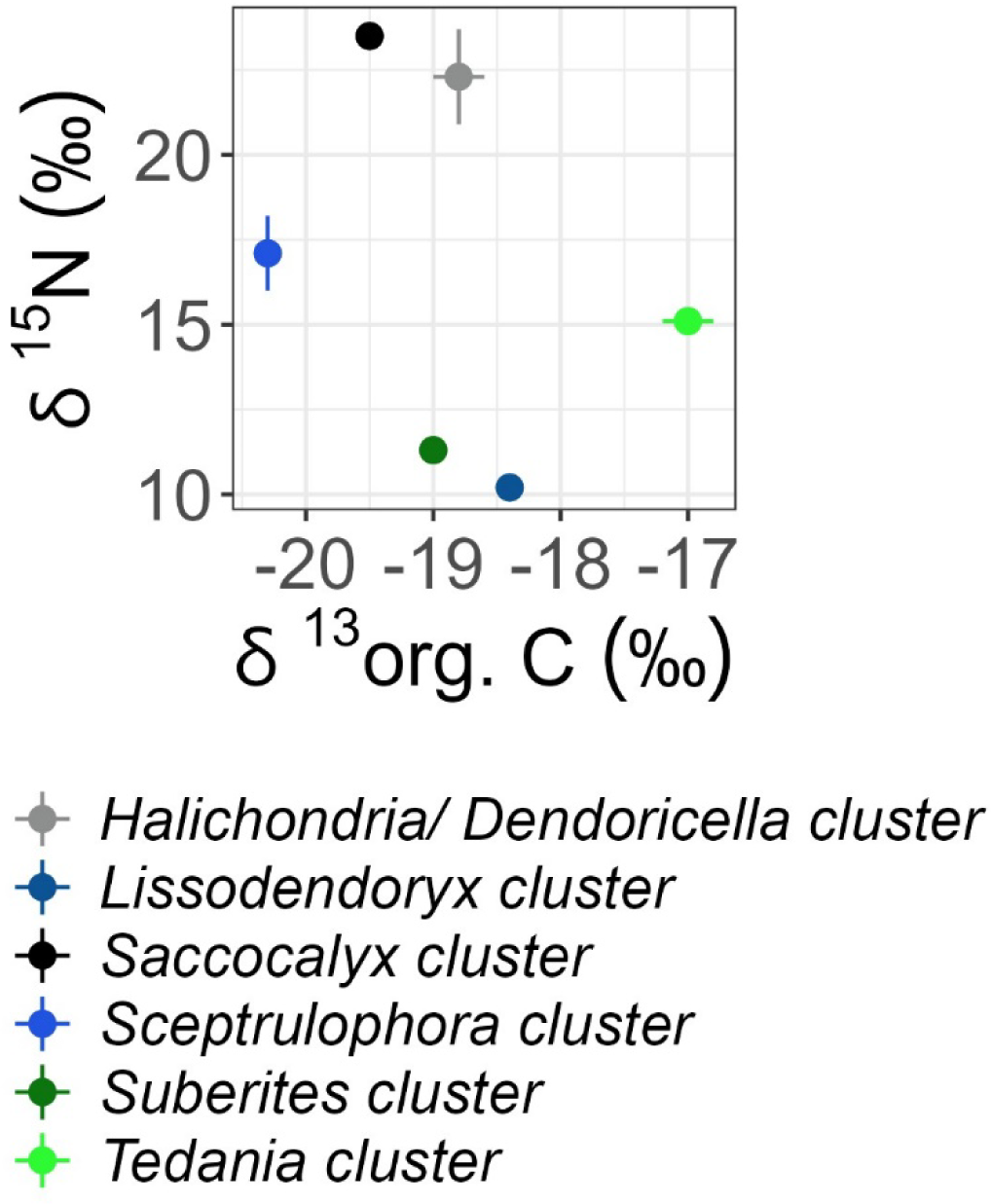
Isotopic composition of carbon (δ^13^org. C, ‰) and nitrogen (δ^15^N, ‰) of sponge tissue collected from New Zealand. Error bars represent 1 standard error.

### 3.2 Nutrient fluxes and oxygen consumption

All sponges released nitrite and ammonium (Fig. 5), but only the nitrite release of the *Halichondria*/ *Dendoricella* cluster (1.45 ± 0.35 µmol g C^−1^ d^−1^) and the *Suberites* cluster (2.17 ± 0.44 µmol g C^−1^ d^−1^) and the ammonium release of the *Tedania* cluster (2.25 ± 0.55 µmol g C^−1^ d^−1^) and the *Halichondria*/ *Dendoricella* cluster (90.7 ± 22.8 µmol g C^−1^ d^−1^) were significantly different from 0 µmol g C^−1^ d^−1^; though, no statistical analyses of nutrient and oxygen fluxes were performed for the *Saccocalyx* cluster and the *Lissodendoryx* cluster (Fig. 5; Table S4). Nitrate was released by the *Halichondria*/ *Dendoricella* cluster and the *Lissodendoryx* cluster, and consumed by the *Suberites* cluster, the *Tedania* cluster, and the Sceptrulophora cluster, though all statistically tested fluxes were not significantly different from 0 µmol g C^−1^ d^−1^ (Fig. 5; Table S4). Phosphate release was only significantly different from 0 µmol g C^−1^ d^−1^ in the case of the Sceptrulophora cluster (2.10 ± 0.47 µmol g C^−1^ d^−1^) and the *Halichondria*/ *Dendoricella* cluster (18.5 ± 4.83 µmol g C^−1^ d^−1^) (Fig. 5; Table S4). Silicon was released by the *Saccocalyx* cluster and the *Halichondria*/ *Dendoricella* cluster and it was consumed by the *Suberites* cluster, the *Lissodendoryx* cluster, the Sceptrulophora cluster, and the *Tedania* cluster, though none of the statistically tested fluxes was significantly different from 0 µmol g C^−1^ d^−1^ (Fig. 5; Table S4).

**Figure 5.**
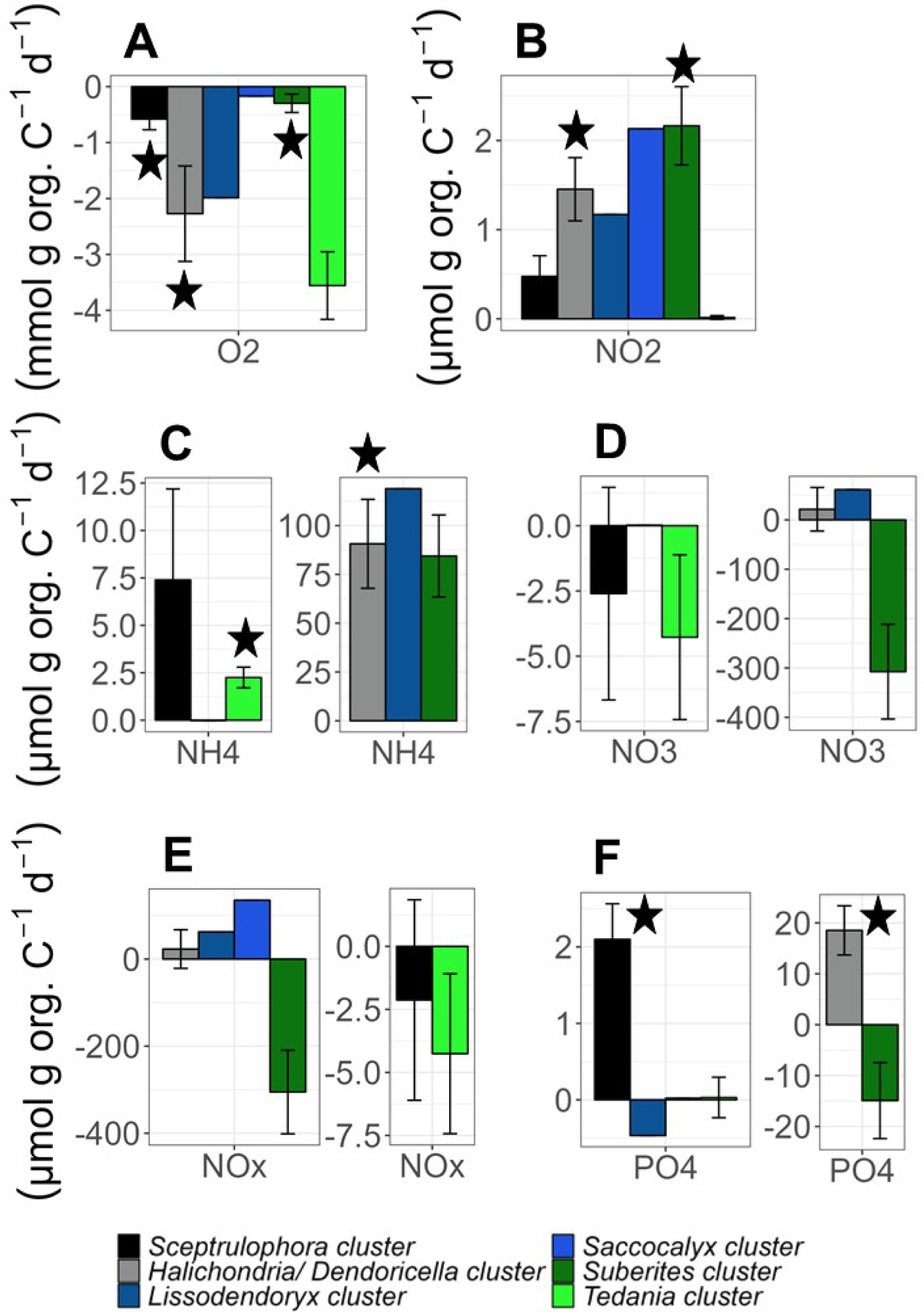
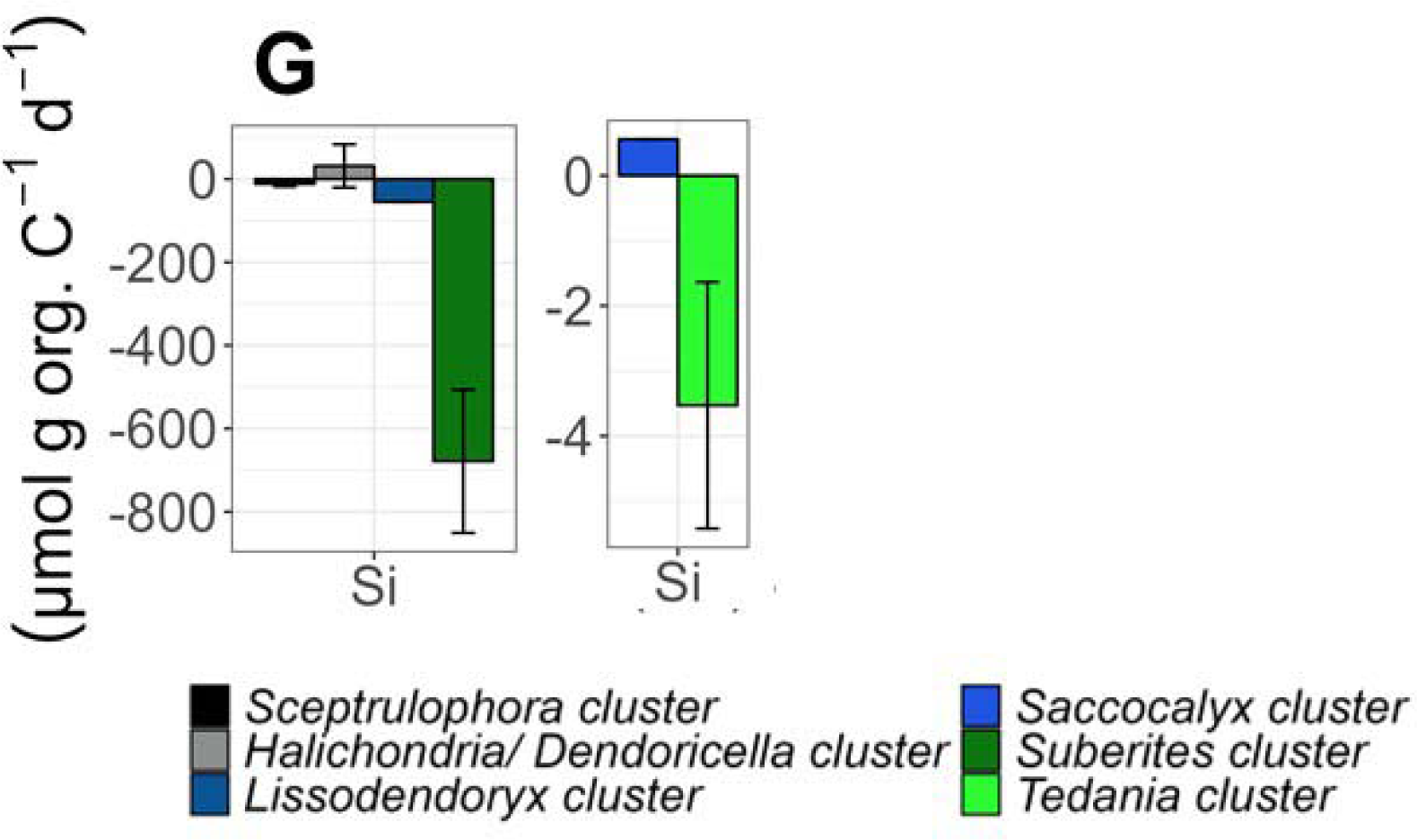
Fluxes (mean ± 1 standard error) of (A) oxygen (O_2_; mmol O_2_ g org. C^1^ d^−1^), (B) nitrite (NO_2_^−^), (C) ammonium (NH_4_^+^), (D) nitrate (NO_3_^−^), (E) NO_x_^−^ (i.e., nitrate + nitrite), (F) phosphate (PO_4_^3-^), and (G) silicon [Si] (all µmol g org. C^−1^ d^−1^). Negative fluxes signify that the sponges took up a specific compound, whereas positive fluxes mean that sponges released this compound. A star symbolizes that the average of this specific flux is significantly different from 0. No statistical analyses were performed for fluxes of the *Saccocalyx* cluster and the *Lissodendoryx* cluster, because only single sponges of these clusters were incubated. A table with nutrient fluxes calculated per sponge volume (cm^3^), sponge wet mass (g *WM*), sponge dry mass (g *DM*), and sponge ash-free dry mass (g *AFDM*) is presented in Table S5.

Sponges consumed between 0.17 mmol O_2_ g C d^−1^ (*Saccocalyx* cluster) and 3.56±0.60 mmol O_2_ g C d^−1^ (*Suberites* cluster) oxygen (Fig. 5), but the oxygen consumption rate of the *Tedania* cluster was not significantly different from 0 mmol O_2_ g C d^−1^ (Table S4). Oxygen fluxes presented per sponge volume, sponge wet/ dry/ ash-free dry mass are presented in Table S5.

### 3.3 Fatty acid composition of sponges

The PLFA compositions (Fig. 6A, B; Table S6) differed strongly among sponge clusters. Between 24.2% (*Halichondria*/ *Dendoricella* cluster) and 59.4 % (*Saccocalyx* cluster) of the PLFAs found in sponges consisted of long-chain fatty acids (LCFA, i.e., fatty acid with ≥24 C atoms) (Fig. 6A). The other PLFA classes detected in sponges were branched fatty acids (0.84 ± 0.57%), cyclic fatty acids (2.36 ± 1.67%), highly unsaturated fatty acids (HUFA, i.e., fatty acids with 4 double bonds; 3.61 ± 1.50%), methyl-fatty acids (0.73 ± 0.38%), monosaturated fatty acids (MUFA; 11.3 ± 2.14%), polyunsaturated fatty acids (PUFAs, i.e., fatty acids with ≥2 double bonds; 0.35 ± 0.35%), saturated fatty acids (SFA; 23.8 ± 2.77%), and undefined fatty acids (20.4 ± 3.03%).

**Figure 6.**
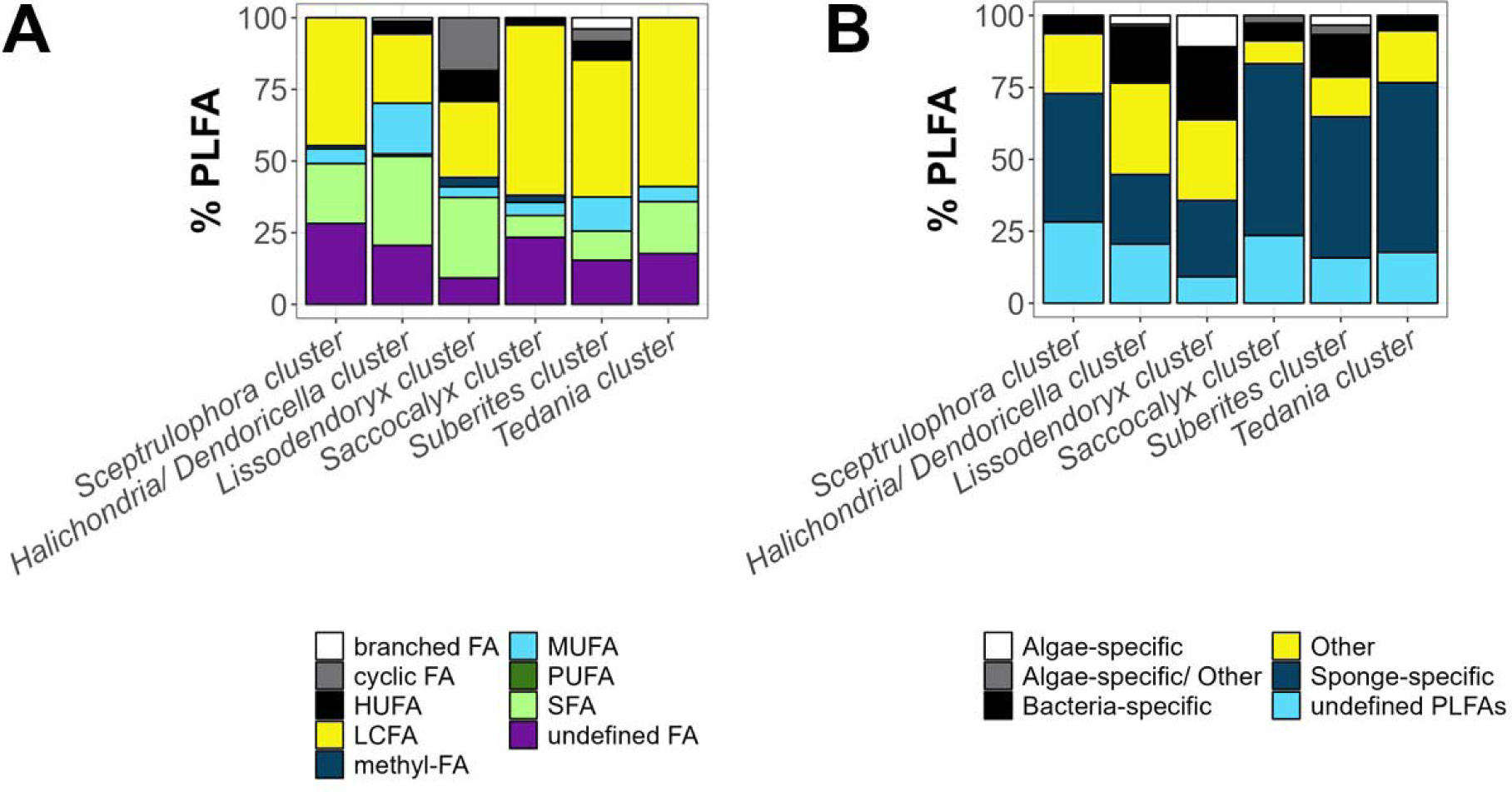
(**A**) Contribution (%) of individual phospholipid-derived fatty acid (PLFA) classes to the total concentrations in sponges. (**B**) Contribution (%) of individual PLFA categories to total PLFA concentrations in sponges. Abbreviations: branched FA = branched fatty acid, HUFA = highly unsaturated fatty acid, LCFA = long-chain fatty acid, methyl-FA = methyl-fatty acid, MUFA = monounsaturated fatty acid, PUFA = polyunsaturated fatty acid, SFA = saturated fatty acid.

The highest concentrations of PLFAs isolated from the sponge tissue were sponge-specific PLFAs (min: 24.2% of total PLFA concentration, *Halichondria*/ *Dendoricella* cluster; max: 59.7%, *Saccocalyx* cluster) (Fig. 6B). The PLFA categories with the second and third highest concentrations were others (min: 7.95% *Saccocalyx* cluster; max: 31.9% *Halichondria*/ *Dendoricella* cluster) and bacteria-specific PLFAs (min: 5.25%, *Tedania* cluster; max: 25.3%, *Lissodendoryx* cluster). About 9.21% (*Lissodendoryx* cluster) to 28.2% (Sceptrulophora cluster) of the total PLFA concentration consisted of undefined PLFAs.

### 3.4 Composition of sponge-associated bacterial community

The bacterial community in all sponge clusters, except for the *Suberites* cluster, were dominated by Proteobacteria (min: 84.8±5.58% ASVs *Tedania* cluster; max: 95.3±0.82% ASVs *Halichondria*/ *Dendoricella* cluster), followed by Bacteroidota (0.28±0.01% − 5.50±2.67% ASVs), Planctomycetota (0.26±0.08% − 1.11±0.44% ASVs), and Nitrospinota (0.03±0.02% − 3.34±1.74% ASVs) (Fig. 7, Table S7). The most abundant Bacteria in the *Suberites* cluster were Proteobacteria (44.3% ASVs), Chloroflexota (20.3% ASVs), Acidobacteriota (13.7% ASVs), and Actinobacteriota (3.10% ASVs) (Fig. 7).

**Figure 7.**
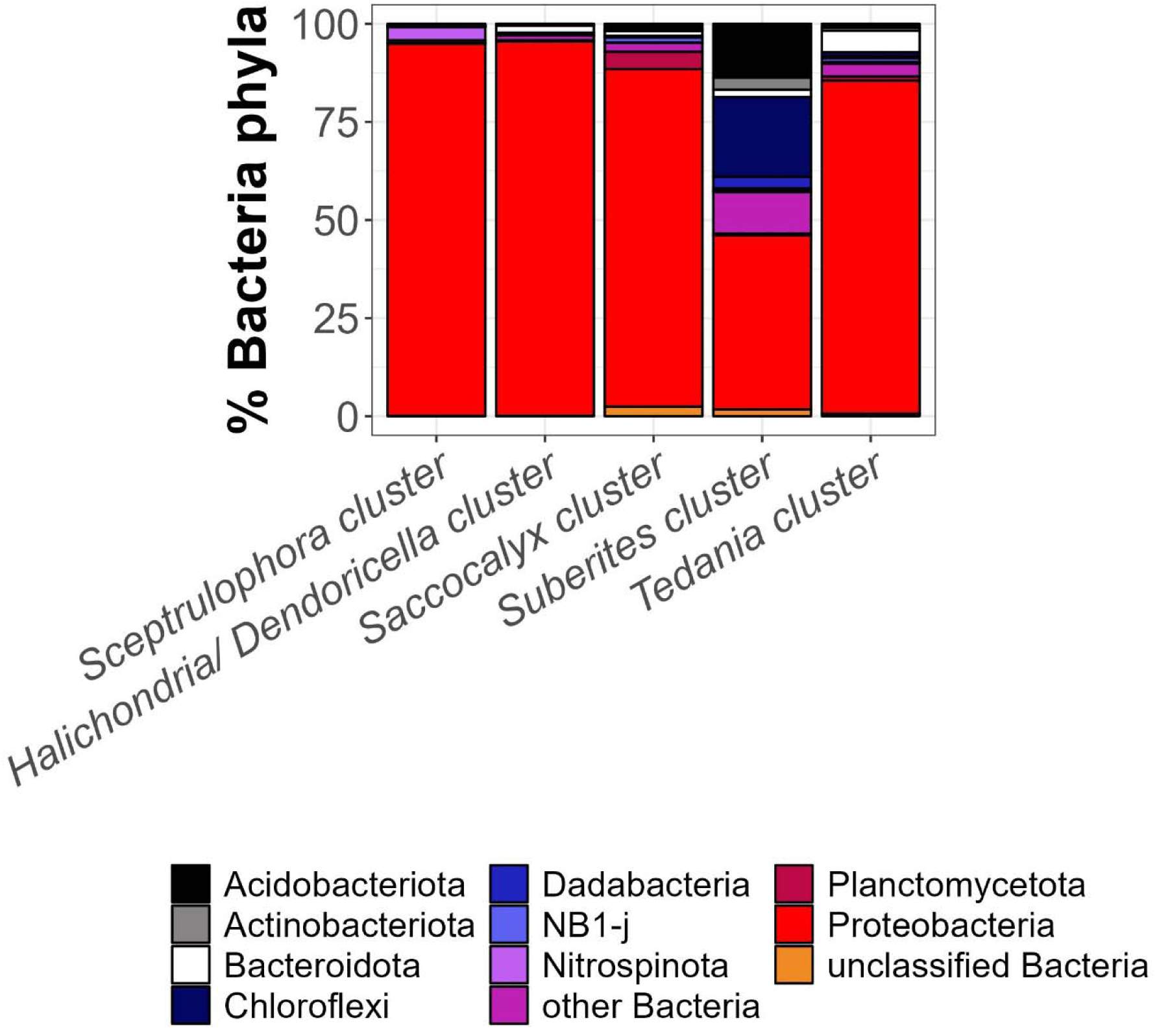
Relative abundance of bacteria phyla (%) in the five analyzed sponge clusters. The nine most abundant microbial phyla are indicated by names, others are aggregated under the category “other Bacteria” and “unclassified Bacteria”.

Statistical comparisons revealed no significant differences in bacterial community composition between the Sceptrulophora cluster, *Tedania* cluster, or *Halichondria*/ *Dendoricella* cluster (Table 2).

**Table 2.**
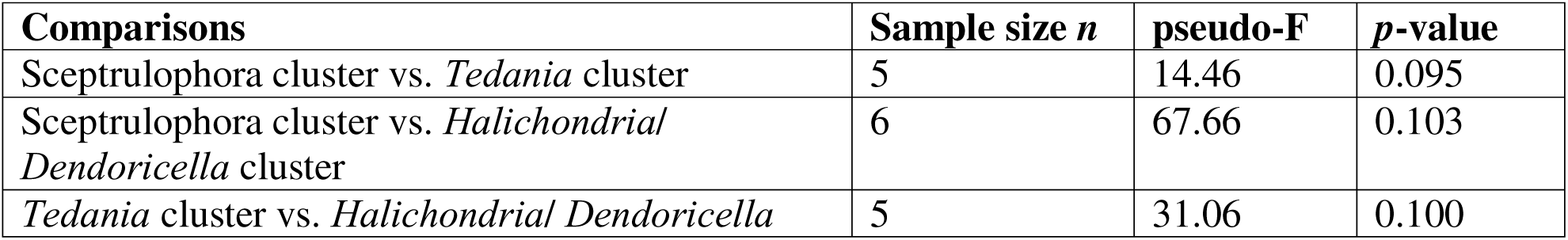
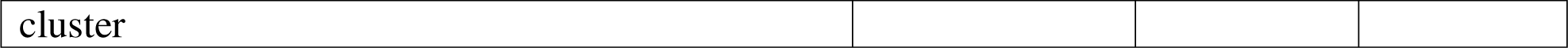
Results of pairwise PERMANOVAs to assess statistical differences in bacterial community composition between sponge clusters.

## 4. Discussion

### 4.1 Nitrogen fluxes of deep-sea sponges

The dissolved inorganic nitrogen (DIN) dynamics of the sponge clusters collected from New Zealand comprised the uptake of nitrate and the release of ammonium, nitrate, and nitrite. Most ammonium was excreted by the *Lissodendoryx* cluster, whereas most NO_x_^−^ (i.e., nitrate + nitrite) release was detected in the *Suberites* cluster. A high excretion of NO_x_^−^ by the HMA sponge *Suberites* spp. indet. is not uncommon and has been observed in other studies of HMA sponges from Florida Bay (W Atlantic) (Hoer et al., 2018; Southwell et al., 2008), whereas LMA sponges, such as all other sponge clusters investigated in this study, predominantly excrete ammonium (Hoer et al., 2018).

A more detailed look at the DIN fluxes measured *ex situ* and *in situ* revealed three different types of nitrogen cycling in sponges. Type 1 is represented by the release of ammonium and NO_x_^−^ by *Dendoricella* spp. indet. and the *Lissodendoryx* cluster (Fig. S4). Type 2 is the release of ammonium and the uptake of NO_x_^−^ by the Sceptrulophora cluster, *Halichondria* sp. indet., the *Suberites* cluster, and the *Tedania* cluster and type 3 comprises the uptake of ammonium and the release of NO_x_^−^ by the *Saccocalyx* cluster.

During type 1, the sponge holobiont aerobically respires and ammonificates organic matter to ammonium, fixes N_2_ to ammonium (likely a minor process), and aerobically nitrifies heterotrophically produced ammonium to nitrate and nitrite. This type of N cycling is not limited to the investigated deep-sea sponges from New Zealand, but has also been observed in the shallow-water sponges *Ircinia felix* (Duchassaing & Michelotti, 1864), *Ircinia campana* (Lamarck, 1814), and *Dasycladus* sp. (previously *Spongia* sp.) from Florida Bay and *Xestospongia muta* (Schmidt, 1870) from the Caribbean (Fiore et al., 2013; Hoer et al., 2018). In the investigated sponges from New Zealand, however, the dominating N cycling processes might vary as the δ^15^N values of sponge tissue differ substantially between *Dendoricella* spp. indet. and the *Lissodendoryx* cluster. The very high δ^15^N *Dendoricella* spp. indet. tissue value (23.8±2.36‰) resembles nitrification which is believed to lead to a net enrichment in δ^15^N of sponge tissue (Southwell, 2007). In comparison, the lower δ^15^N *Lissodendoryx* cluster tissue value (10.2‰) indicates that this cluster nitrifies, but also likely feeds more selectively than *Dendoricella* spp. indet. and/ or takes up more dissolved organic matter (DOM) and/ or fixes more N_2_ than *Dendoricella* spp. indet. (Southwell, 2007). Unfortunately, we did not take samples for bacterioplankton, DOM or dissolved N_2_ concentrations to verify these hypotheses.

During type 2, the sponge holobiont aerobically respires and ammonificates organic matter to ammonium and reduces nitrate anaerobically to ammonium via dissimilatory nitrate reduction to ammonium (DNRA). Dissimilatory nitrate reduction to ammonium requires anaerobic niches or ‘pockets’ within the sponge holobiont and, indeed, Kumala et al., (2021) measured oxygen gradients inside the temperate shallow-water sponge *Halichondria (Halichondria) panicea* (Pallas, 1766) and Gatti et al., (2002) detected differences in oxygen saturation inside sponge tissue of the Antarctic sponge *Suberites domuncula* (Olivi, 1792). In these anaerobic niches, DNRA is driven by the microbiome of the sponge holobiont, such as Planctomycetota and Actinobacteriota that are involved in DNRA in the deep-sea sponge *Vazella pourtalesii* (Schmidt, 1870) from Emerald Basin (NW Atlantic) (Maldonado et al., 2021).

Type 3 resembles the microbial nitrification of ammonium to nitrite and subsequently to nitrate by ammonium-oxidizing Bacteria (AOB) and/ or Archaea (AOA) (Hallam et al., 2006; Hentschel et al., 2006; Preston et al., 1996; Steinert et al., 2020). Like the *Saccocalyx* cluster in this study, *Nodastrella nodastrella* (Topsent, 1915) from Rockall Bank (NE Atlantic) and *V. pourtalesii* from Emerald Basin take up ammonium (Maldonado et al., 2021; Van Duyl et al., 2008) and mostly release nitrite instead of nitrate indicating that a subsequent oxidation of nitrite to nitrate does not happen within the sponge holobiont, potentially due to a lack of nitrite oxidizing bacteria (NOB) (Maldonado et al., 2021).

### 4.2 Silicon fluxes of deep-sea sponges

The uptake of silicon by the sponges from New Zealand varied substantially and ranged from silicon release by the *Saccocalyx* cluster and by *Dendoricella* spp. indet. of the *Halichondria*/ *Dendoricella* cluster to the uptake of silicon by the other sponge clusters in the order of 10^−2^ to 10 μmol Si g^−1^ *AFDM* sponge h^−1^. This large difference in silicon fluxes might be the result of (1) species-specific differences in silicon uptake (Perea-Blázquez et al., 2012), (2) reduced silicon uptake due to starvation (Frøhlich and Barthel, 1997) or (3) during reproduction (Jones, 1987) in (Jones, 1991) (Frøhlich and Barthel, 1997), and/or (4) low/ limiting ambient silicon concentrations (Frøhlich and Barthel, 1997; Reincke and Barthel, 1997). As we know too little about the natural history and life cycle of the investigated deep-sea sponge clusters from New Zealand, we cannot pinpoint which of the processes is responsible for the differences in silicon uptake, however we try to identify the most likely process for few of the sponge clusters.

Lüskow et al., (2019) interpreted a low sponge *CI* value (mean ± standard deviation: 0.53 ± 0.09) in winter for *H. panicea* as a sign of starvation. All New Zealand sponges in this study had sponge CI values that were lower than the low winter CI of *H. panicea*, implying that they were likely food limited. The *Suberites* cluster, though, with its highest silicon uptake rate of 678 ± 172 µmol g org. C^−1^ d^−1^ was either not starving or the food limitation was minimal for this species. Unfortunately, we did not collect any *in-situ* water samples of dissolved organic matter, particulate organic matter, or bacterioplankton concentrations to explore this further.

In a study of *H. panicea* from the Kiel Bight (Baltic Sea, Germany), Frøhlich and Barthel (1997) measured the highest silicon uptake in specimens that showed no stages of late oogenesis and/ or embryogenesis. However, we did not investigate reproductive stages and therefore cannot provide any information in favor or against the hypothesis that silicon fluxes in the investigated sponge clusters were influenced by a specific stage of the reproductive cycle.

The uptake of silicon by sponges follows saturable Michaelis-Menten kinetics (López-Acosta et al., 2018; Reincke and Barthel, 1997) and increases with increasing ambient silicon concentrations until the limiting silicon uptake rate is reached. As we incubated each sponge cluster at similar silicon concentrations (total range across all incubations: 4.58 − 69.8 µmol l^−1^ Si), we cannot determine at which point of the Michaelis-Menten curve the silicon uptake happened and therefore we cannot identify whether sponges were silicon limited. In fact, experimental data from incubations of *Haliclona (Haliclona) simulans* (Johnston, 1842) and *Suberites ficus* (Johnston, 1842) found saturated uptake rates at 70 µmol l^−1^ Si (*H. simulans*) and 130 µmol l^−1^ Si (*S. ficus*) (López-Acosta et al., 2018), suggesting that all sponge specimens investigated in this study were silicon limited, except for the specimen of the *Saccocalyx* cluster. This specimen, however, released most silicon which might be related to degradation processes of the sponge and confound the measured ambient silicon concentration.

### 4.3 Bacterial community of the sponge holobionts

Microbial fingerprints indicated that all sponge clusters, except the *Suberites* cluster, were likely LMA sponges, being dominated by Proteobacteria. The *Suberites* cluster, however, was likely a HMA sponge, indicated among others by a large fraction of Chloroflexota bacteria and Acidobacteriota, besides Proteobacteria. Overall, the investigated sponges hosted microbiomes whose bacterial composition was comparable to other deep-sea sponges (Acidobacteriota, Chloroflexota, and Dadabacteria enriched in HMA over LMA sponges; Bacteroidota and Proteobacteria enriched in LMA over HMA), and resembled similar patterns as described in another study on deep-sea sponges from the same research cruise around New Zealand (Steinert et al., 2020).

In *V. pourtalesii* Proteobacteria are involved in the assimilation of ammonium, ammonification, and DOM processing and some Proteobacteria species contain the enzyme ammonia-monooxygenase (AMO) which is responsible for the oxidation of ammonium to hydroxylamine, the first step of the nitrification (Maldonado et al., 2021). Hence, Proteobacteria present in *Dendoricella* spp. indet. and *Lissodendoryx* cluster might be involved in the ammonification of DOM. These bacteria could also participate in denitrification as clusters of Beta- and Gammaproteobacteria detected in *G. barretti* have the gene *nirS* encoding for the enzyme cytochrome cd1-containing nitrite reductase (Hoffmann et al., 2009). However, as we neither measured nitric oxide, not did we perform metatranscriptomics of the microbiome in our sponge clusters, we cannot confirm this.

Bacteroidota play a key role in tropical coral reefs, where they are involved in amino acid metabolism and biosynthesis of terpenoids and polyketides in seawater (Glasl et al., 2020). They can contain *SusD*-like genes and genes encoding for glycoside hydrolase pointing towards the ability of degrading polysaccharides (Glasl et al., 2020; Mackenzie et al., 2012). Indeed, Bacteroidota present in the shallow-water coral reef sponges *Ircinia ramosa* (Keller, 1889), *Ircinia microconulosa* Pulitzer-Finali, 1982 and *Phyllospongia foliascens* (Pallas, 1766) are involved in carbohydrate degradation (O’Brien et al., 2023). Also in *V. pourtalesii*, they contribute to DOM processing (Maldonado et al., 2021). Hence, Bacteroidota enriched in the *Tedania* cluster compared to other sponge clusters investigated here might be involved in the degradation of carbohydrates and ammonification of DOM which was measured as ammonium release.

Planctomycetota are involved in the nitrogen cycle of the sponge holobiont. In fact, metatranscriptomics of *V. pourtalesii* showed that members of this bacteria phylum have genes encoding for nitrate reductase which is necessary for the denitrification of nitrate to nitrite (Maldonado et al., 2021). Furthermore, they contain the genes *nirB/ D* that encode for NADH dependent nitrate reductase that is part of the DNRA pathway and they accommodate genes for nitrogenase enzymes required for N_2_ fixation to ammonium (Maldonado et al., 2021). In this study, however, Planctomycetota were most abundant in the *Saccocalyx* cluster where inorganic nutrient flux data indicate that neither DNRA nor denitrification happened at higher rates. We therefore expect that Archaea which were not investigated in this study might have been more active in nitrogen cycling of the *Saccocalyx* cluster than Planctomycetota.

Members of the phylum Nitrospinota are NOB that oxidize nitrite to nitrate and play an important role in dark carbon fixation (Pachiadaki et al., 2017) and nitrite oxidation in oxygen minimum zones (Sun et al., 2019). These bacteria are present in the shallow-water sponges *I. ramosa* and *Coscinoderma mathewsi* (Lendenfeld, 1886) (Australian Great Barrier Reef; (Engelberts et al., 2020; Glasl et al., 2020)), *Petrosia* (*Petrosia*) *ficiformis* (Poiret, 1789) (Eastern Mediterranean Sea; (Burgsdorf et al., 2022)), and *Aplysina aerophoba* (Nardo, 1833) (Adriatic Sea; (Burgsdorf et al., 2022)). In this study, they were most abundant in the *Farrea* cluster (3.34±1.74% ASVs, Table S7), but as we measured (not significant) release of nitrite instead of uptake (Fig. 5), Nitrospinota were likely not very active in this sponge cluster.

Chloroflexota, which was the second most abundant bacteria phylum in the *Suberites* cluster, often dominate the bacterial community of HMA sponge holobionts (Bayer et al., 2018; Busch et al., 2020; Moitinho-Silva et al., 2017). Members of this phylum are aerobic heterotrophs that potentially perform autotrophic carbon fixation in parts of the sponge tissue that becomes anoxic when the sponge pumping activity ceases (Bayer et al., 2018). They also contain genes for fatty acid biosynthesis and degradation (Bayer et al., 2018). Two Chloroflexota classes (i.e., *Anaerolineae* and *Caldilineae*), in particular, are capable of extensive uptake of carbohydrates and their subsequent degradation (Bayer et al., 2018) and it has been speculated that Chloroflexota bacteria are even able to degrade recalcitrant DOM (Bayer et al., 2018; Landry et al., 2017). Hence, we hypothesize that the *Suberites* cluster might take up DOM, though we did not measure DOM fluxes in the incubations and therefore cannot confirm this hypothesis.

### 4.4 Fatty acid composition of deep-sea sponges

Knowledge about differences in (phospholipid-derived) fatty acid profiles of demosponges compared to glass sponges is still very limited, but the data available so far indicate that demosponges synthesize mainly dienoic long-chain fatty acids with 24 to 28 C-atoms, whereas glass sponges have long-chain fatty acids with mostly C30-polyunsaturated fatty acids (Thiel et al., 2002). Furthermore, demosponges contain methyl-branched carbon chains that are less common in glass sponges and brominated long-chain fatty acids which have not been observed in glass sponges (Garson et al., 1994; Thiel et al., 2002). The fatty acid composition of the sponges that were investigated here mostly confirm the previous observations: The concentration of long-chain fatty acids with 24 to 27 C-atoms was higher in demosponges than in glass sponges (Fig. S5). Fatty acids with 28 to 30 C-atoms, however, had higher concentrations in glass sponges than in demosponges (Fig. S5).

A closer look at the fatty acid composition of the different sponge clusters may provide additional information about food sources of these sponges. The presence of mid-chain branched fatty acids (8/9/10/11-Me-C16:0, 9/10/11-Me-C18) that are bacteria-originating precursor fatty acids for long-chain fatty acids (de Kluijver et al., 2021) show that the deep-sea sponge clusters *Farrea*, *Saccocalyx*, and *Lissodendoryx* gain metabolic energy from their endosymbionts, similar to what has been described for Geodiidae sponges from the Northeast Atlantic (de Kluijver et al., 2021).

The presence of algae-specific fatty acids in the *Lissodendoryx* cluster, the *Halichondria*/ *Dendoricella* cluster and the *Suberites* cluster suggests that these sponges take up phytodetritus-derived POC. The results of the DIN flux measurements of the *Lissodendoryx* cluster and the *Suberites* cluster are evidence for this suggestion because both sponge clusters likely ammonificate organic matter. However, the algae-specific fatty acid C20:5ω3 (eicosapentaenoic acid *EPA*) can also be synthesized by deep-water corals, like *Desmophyllum pertusum* (Linnaeus, 1758) (Mueller et al., 2014). Hence, the presence of *EPA* which co-eluted with C20:4ω6 (α-linolenic acid *ARA*) indicates that the *Saccocalyx* cluster, the *Halichondria*/ *Dendoricella* cluster, and the *Suberites* cluster might also consume DOM originating from the mucus of deep-water corals. In fact, ROV imagery from the sampling stations show the presence of deep-water corals in the vicinity of the *Suberites* cluster (Fig. S6).

## 5. Conclusions

In this study, we assessed the role of deep-water sponge holobionts from New Zealand in nitrogen cycling and tried to decipher the potential food-source preferences of these sponges. Sponges of the LMA Sceptrulophora cluster potentially gain metabolic energy from their symbionts and host AOB that might oxidize ammonia to nitrite. Members of the LMA *Saccocalyx* cluster might gain metabolic energy from endosymbionts, such as Proteobacteria, and might host bacteria that nitrify ammonium to nitrite. *Dendoricella* spp. indet. of the *Halichondria*/ *Dendoricella* cluster host Proteobacteria that might contribute to the ammonification of DOM and *Halichondria* sp. of the same cluster potentially contain ammonia-oxidizing or nitrifying bacteria involved in DNRA and ammonification of DOM. Sponges of the *Lissodendoryx* cluster are LMA sponges that might feed selectively on phytodetritus-derived POC which is subsequently ammonificated, but they might also gain metabolic energy from their symbionts. The last investigated LMA sponges belong to the *Tedania* cluster that hosts Bacteroidota which might be able to degrade carbohydrates and contribute to the ammonification of DOM. Additionally, the sponges might contain nitrifying bacteria. In contrast, the *Suberites* cluster sponges are HMA sponges and potentially consume carbohydrates originating from deep-sea coral mucus that are degraded by Chloroflexota. Furthermore, they might ammonificate phytodetritus-based POC. These proposed processes should be confirmed with further experimental studies such as DNA-stable isotope probing (DNA-SIP; e.g., (Campana et al., 2021)) combined with compound-specific stable isotope analysis (CSIA), and metagenomic data.

## CRediT Roles

**Tanja Stratmann**: Conceptualization, Data curation, Formal Analysis, Funding acquisition, Investigation, Methodology, Project administration, Resources, Software, Validation, Visualization, Writing – original draft; **Kathrin Busch**: Data curation, Formal Analysis, Investigation, Methodology, Software, Validation, Visualization, Writing – review & editing; **Anna de Kluijver**: Investigation, Methodology, Writing – review & editing ; **Michelle Kelly**: Taxonomic identification, Writing – review & editing; **Sadie Mills**: Data curation, Resources, Validation, Writing – review & editing; **Sven Rossel**: Investigation, Methodology, Validation, Writing – review & editing; **Peter Schupp**: Funding acquisition, Resources, Supervision, Writing – review & editing.

## Data availability

All data presented in this manuscript are available at Pangaea (**XXX**) and on GenBank (accession numbers of sponge data are presented in Table S1; accession number of bacterial data: **XX**). Molecular genetic data are deposited in the barcode of life data system (BOLD) in the project Sponges off New Zealand (SONZ) and processed mass spectrometry data can be found in Supplementary Data S1.

Sample collection was carried out under the “Application for consent to conduct marine scientific research in areas under national jurisdiction of New Zealand (dated 7.6.2016).”

## Acknowledgements

We greatly acknowledge the crew and scientific party of RV *Sonne* cruise SO254, as well as the ROV Kiel 6000 team from GEOMAR (Kiel) for their valuable support at sea. We also thank Sven Rohde, Tessa Clemens, and the entire benthic invertebrate team of the RV *Sonne* cruise SO254 for their assistance in sample collection, processing, and cataloging. We further acknowledge the analytical support from Peter van Breugel and Jan Peene (NIOZ) and Klaas Nierop (Utrecht University).

This research received funding from the Royal Netherlands Academy of Arts and Sciences (KNAW, The Netherlands) to TS (Academy Ecology Grant 2017) and from the Dutch Research Council (NWO, The Netherlands) to TS (NWO-Rubicon grant no. 019.182EN.012, NWO-Talent program Veni grant no. VI.Veni.212.211). AdK was supported by the SpongGES project of the European Union’s Horizon 2020 research and innovation program (grant no. 679849). PS received funding by the Federal Ministry of Education and Research (BMBF) for the cruise SO254, grant no. 03G0254A, PORIBACNEWZ.

Specimens were collected as part of the project “PoriBacNewZ” by the Institut für Chemie und Biologie des Meeres (ICBM), University Oldenburg on the German flagship RV *Sonne*, using the ROV Kiel 6000 (GEOMAR) with participation and funding from GEOMAR, DSMZ, LMU, NIOZ, NIWA, and ETH-Zurich. NIWA voyage participation was funded through the MBIE SSIF Enhancing Collections project.

This is publication number 21 of Senckenberg am Meer Proteome Laboratory and number 96 of Senckenberg am Meer Metabarcoding and Molecular Laboratory.

## References

Altschul, S.F., Madden, T.L., Schäffer, A.A., Zhang, J., Zhang, Z., Miller, W., Lipman, D.J., 1997. Gapped BLAST and PSI-BLAST: A new generation of protein database search programs. Nucleic Acids Res 25, 3389–3402.

Bart, M.C., de Kluijver, A., Hoetjes, S., Absalah, S., Mueller, B., Kenchington, E., Rapp, H.T., de Goeij, J.M., 2020. Differential processing of dissolved and particulate organic matter by deep-sea sponges and their microbial symbionts. Sci Rep 10, 17515. 10.1038/s41598-020-74670-0

Bayer, K., Jahn, M.T., Slaby, B.M., Moitinho-Silva, L., Hentschel, U., 2018. Marine sponges as Chloroflexi hot spots: Genomic insights and high-resolution visualization of an abundant and diverse symbiotic clade. mSystems 3. 10.1128/mSystems.00150-18

Beaulieu, S.E., 2001. Life on glass houses: Sponge stalk communities in the deep sea. Mar Biol 138, 803–817. 10.1007/s002270000500

Bell, J.J., 2008. The functional roles of marine sponges. Estuar Coast Shelf Sci 79, 341–353. 10.1016/j.ecss.2008.05.002

Bligh, E.L.G., Dyer, W.J.A., 1959. A Rapid Method of Total Lipid Extraction And Purification. Can J Biochem Physiol 37, 911–917. 10.1139/o59-099

Bolyen, E., Rideout, J.R., Dillon, M.R., Bokulich, N.A., Abnet, C.C., Al-Ghalith, G.A., Alexander, H., Alm, E.J., Arumugam, M., Asnicar, F., Bai, Y., Bisanz, J.E., Bittinger, K., Brejnrod, A., Brislawn, C.J., Brown, C.T., Callahan, B.J., Caraballo-Rodríguez, A.M., Chase, J., Cope, E.K., Da Silva, R., Diener, C., Dorrestein, P.C., Douglas, G.M., Durall, D.M., Duvallet, C., Edwardson, C.F., Ernst, M., Estaki, M., Fouquier, J., Gauglitz, J.M., Gibbons, S.M., Gibson, D.L., Gonzalez, A., Gorlick, K., Guo, J., Hillmann, B., Holmes, S., Holste, H., Huttenhower, C., Huttley, G.A., Janssen, S., Jarmusch, A.K., Jiang, L., Kaehler, B.D., Kang, K. Bin, Keefe, C.R., Keim, P., Kelley, S.T., Knights, D., Koester, I., Kosciolek, T., Kreps, J., Langille, M.G.I., Lee, J., Ley, R., Liu, Y.-X., Loftfield, E., Lozupone, C., Maher, M., Marotz, C., Martin, B.D., McDonald, D., McIver, L.J., Melnik, A. V., Metcalf, J.L., Morgan, S.C., Morton, J.T., Naimey, A.T., Navas-Molina, J.A., Nothias, L.F., Orchanian, S.B., Pearson, T., Peoples, S.L., Petras, D., Preuss, M.L., Pruesse, E., Rasmussen, L.B., Rivers, A., Robeson, M.S., Rosenthal, P., Segata, N., Shaffer, M., Shiffer, A., Sinha, R., Song, S.J., Spear, J.R., Swafford, A.D., Thompson, L.R., Torres, P.J., Trinh, P., Tripathi, A., Turnbaugh, P.J., Ul-Hasan, S., van der Hooft, J.J.J., Vargas, F., Vázquez-Baeza, Y., Vogtmann, E., von Hippel, M., Walters, W., Wan, Y., Wang, M., Warren, J., Weber, K.C., Williamson, C.H.D., Willis, A.D., Xu, Z.Z., Zaneveld, J.R., Zhang, Y., Zhu, Q., Knight, R., Caporaso, J.G., 2019. Reproducible, interactive, scalable and extensible microbiome data science using QIIME 2. Nat Biotechnol 37, 852–857. 10.1038/s41587-019-0209-9

Burgsdorf, I., Sizikov, S., Squatrito, V., Britstein, M., Slaby, B.M., Cerrano, C., Handley, K.M., Steindler, L., 2022. Lineage-specific energy and carbon metabolism of sponge symbionts and contributions to the host carbon pool. ISME Journal 16, 1163–1175. 10.1038/s41396-021-01165-9

Busch, K., Slaby, B.M., Bach, W., Boetius, A., Clefsen, I., Colaço, A., Creemers, M., Cristobo, J., Federwisch, L., Franke, A., Gavriilidou, A., Hethke, A., Kenchington, E., Mienis, F., Mills, S., Riesgo, A., Ríos, P., Roberts, E.M., Sipkema, D., Pita, L., Schupp, P.J., Xavier, J., Rapp, H.T., Hentschel, U., 2022. Biodiversity, environmental drivers, and sustainability of the global deep-sea sponge microbiome. Nat Commun 13. 10.1038/s41467-022-32684-4

Busch, K., Wurz, E., Rapp, H.T., Bayer, K., Franke, A., Hentschel, U., 2020. Chloroflexi Dominate the Deep-Sea Golf Ball Sponges Craniella zetlandica and Craniella infrequens Throughout Different Life Stages. Front Mar Sci 7. 10.3389/fmars.2020.00674

Callahan, B.J., McMurdie, P.J., Rosen, M.J., Han, A.W., Johnson, A.J.A., Holmes, S.P., 2016. DADA2: High-resolution sample inference from Illumina amplicon data. Nat Methods 13, 581–583. 10.1038/nmeth.3869

Campana, S., Busch, K., Hentschel, U., Muyzer, G., de Goeij, J.M., 2021. DNA-stable isotope probing (DNA-SIP) identifies marine sponge-associated bacteria actively utilizing dissolved organic matter (DOM). Environ Microbiol 23, 4489–4504. 10.1111/1462-2920.15642

Caporaso, J.G., Lauber, C.L., Walters, W.A., Berg-Lyons, D., Lozupone, C.A., Turnbaugh, P.J., Fierer, N., Knight, R., 2011. Global patterns of 16S rRNA diversity at a depth of millions of sequences per sample. Proc Natl Acad Sci U S A 108, 4516–4522. 10.1073/pnas.1000080107

Carballeira, N., Thompson, J.E., Ayanoglu, E., Djerassi, C., 1986. Biosynthetic studies of marine lipids. 5. The biosynthesis of long-chain branched fatty acids in marine sponges. J Org Chem 51, 2751–2756. 10.1021/jo00364a024

Cathalot, C., Van Oevelen, D., Cox, T.J.S., Kutti, T., Lavaleye, M., Duineveld, G.C.A., Meysman, F.J.R., 2015. Cold-water coral reefs and adjacent sponge grounds: Hotspots of benthic respiration and organic carbon cycling in the deep sea. Front Mar Sci 2, 1–12. 10.3389/fmars.2015.00037

Conway, K.W., Barrie, J.V., Austin, W.C., Luternauer, J.L., 1991. Holocene sponge bioherms on the western Canadian continental shelf. Cont Shelf Res 11, 771–790. 10.1016/0278-4343(91)90079-L

Conway, K.W., Barrie, J.V., Krautter, M., 2005. Geomorphology of unique reefs on the western Canadian shelf: Sponge reefs mapped by multibeam bathymetry. Geo-Marine Letters 25, 205–213. 10.1007/s00367-004-0204-z

Cornway, K.W., Krautter, M., Barrie, J.V., Neuweiler, M., Conway, K.W., Krautter, M., Barrie, J.V., Neuweiler, M., 2001. Hexactinellid sponge reefs on the Canadian continental shelf: A unique “living fossil.” Geoscience Canada 28, 71–78.

de Kluijver, A., 2021. Fatty acid analysis sponges. protocols.io. 10.17504/protocols.io.bhnpj5dn

de Kluijver, A., Nierop, K.G.J., Morganti, T.M., Bart, M.C., Slaby, B.M., Hanz, U., de Goeij, J.M., Mienis, F., Middelburg, J.J., 2021. Bacterial precursors and unsaturated long-chain fatty acids are biomarkers of North-Atlantic deep-sea demosponges. PLoS One 16, e0241095. 10.1371/journal.pone.0241095

Diaz, C.M., Rützler, K., 2001. Sponges: An essential component of Caribbean coral reefs. Bull Mar Sci 69, 535–546.

Dohrmann, M., Janussen, D., Reitner, J., Collins, A.G., Wörheide, G., 2008. Phylogeny and evolution of glass sponges (Porifera, Hexactinellida). Syst Biol 57, 388–405. 10.1080/10635150802161088

Engelberts, J.P., Robbins, S.J., de Goeij, J.M., Aranda, M., Bell, S.C., Webster, N.S., 2020. Characterization of a sponge microbiome using an integrative genome-centric approach. ISME Journal 14, 1100–1110. 10.1038/s41396-020-0591-9

Erwin, P.M., Coma, R., López-Sendino, P., Serrano, E., Ribes, M., 2015. Stable symbionts across the HMA-LMA dichotomy: Low seasonal and interannual variation in sponge-associated bacteria from taxonomically diverse hosts. FEMS Microbiol Ecol 91. 10.1093/femsec/fiv115

Faergeman, N.J., Knudsen, J., 1997. Role of long-chain fatty acyl-CoA esters in the regulation of metabolism and in cell signalling. Biochemical Journal 323, 1–12.

Feuda, R., Dohrmann, M., Pett, W., Philippe, H., Rota-Stabelli, O., Lartillot, N., Wörheide, G., Pisani, D., 2017. Improved modeling of compositional heterogeneity supports sponges as sister to all other animals. Current Biology 27, 3864–3870. 10.1016/j.cub.2017.11.008

Fiore, C.L., Baker, D.M., Lesser, M.P., 2013. Nitrogen Biogeochemistry in the Caribbean Sponge, Xestospongia muta: A Source or Sink of Dissolved Inorganic Nitrogen? PLoS One 8. 10.1371/journal.pone.0072961

Frøhlich, H., Barthel, · D, 1997. Silica uptake of the marine sponge Halichondria panicea in Kiel Bight. Mar Biol 128, 115–125.

Garson, M.J., Zimmermann, M.P., Battershill, C.N., Holden, J.L., Murphy, P.T., 1994. The distribution of brominated longLchain fatty acids in sponge and symbiont cell types from the tropical marine sponge Amphimedon terpenensis. Lipids 29, 509–516. 10.1007/BF02578249

Gatti, S., Brey, T., Müller, W.E.G., Heilmayer, O., Holst, G., 2002. Oxygen microoptodes: a new tool for oxygen measurements in aquatic animal ecology. Mar Biol 140, 1075–1085. 10.1007/s00227-002-0786-9

Gibb, S., 2015. MALDIquantForeign: Import/Export routines for MALDIquant. A package for R.

Gibb, S., Strimmer, K., 2012. Maldiquant: A versatile R package for the analysis of mass spectrometry data. Bioinformatics 28, 2270–2271. 10.1093/bioinformatics/bts447

Giles, E.C., Kamke, J., Moitinho-Silva, L., Taylor, M.W., Hentschel, U., Ravasi, T., Schmitt, S., 2013. Bacterial community profiles in low microbial abundance sponges. FEMS Microbiol Ecol 83, 232–241. 10.1111/j.1574-6941.2012.01467.x

Glasl, B., Robbins, S., Frade, P.R., Marangon, E., Laffy, P.W., Bourne, D.G., Webster, N.S., 2020. Comparative genome-centric analysis reveals seasonal variation in the function of coral reef microbiomes. ISME Journal 14, 1435–1450. 10.1038/s41396-020-0622-6

Gouy, M., Guindon, S., Gascuel, O., 2010. Sea view version 4: A multiplatform graphical user interface for sequence alignment and phylogenetic tree building. Mol Biol Evol 27, 221–224. 10.1093/molbev/msp259

Graber, S., Sumida, C., Nunez, E., 1994. Fatty acids and cell signal transduction. J Lipid Mediat Cell Signal 9, 91–116.

Hallam, S.J., Mincer, T.J., Schleper, C., Preston, C.M., Roberts, K., Richardson, P.M., DeLong, E.F., 2006. Pathways of carbon assimilation and ammonia oxidation suggested by environmental genomic analyses of marine Crenarchaeota. PLoS Biol 4, 520–536. 10.1371/journal.pbio.0040095

Hentschel, U., Usher, K.M., Taylor, M.W., 2006. Marine sponges as microbial fermenters. FEMS Microbiol Ecol 55, 167–177. 10.1111/j.1574-6941.2005.00046.x

Hildebrand, T., Osterholz, H., Bunse, C., Grotheer, H., Dittmar, T., Schupp, P.J., 2022. Transformation of dissolved organic matter by two Indo-Pacific sponges. Limnol Oceanogr 67, 2483–2496. 10.1002/lno.12214

Hoer, D.R., Tommerdahl, J.P., Lindquist, N.L., Martens, C.S., 2018. Dissolved inorganic nitrogen fluxes from common Florida Bay (U.S.A.) sponges. Limnol Oceanogr 63, 2563–2578. 10.1002/lno.10960

Hoffmann, F., Radax, R., Woebken, D., Holtappels, M., Lavik, G., Rapp, H.T., Schläppy, M.L., Schleper, C., Kuypers, M.M.M., 2009. Complex nitrogen cycling in the sponge Geodia barretti. Environ Microbiol 11, 2228–2243. 10.1111/j.1462-2920.2009.01944.x

Jiménez, E., Ribes, M., 2007. Sponges as a source of dissolved inorganic nitrogen: Nitrification mediated by temperate sponges. Limnol Oceanogr 52, 948–958. 10.4319/lo.2007.52.3.0948

Jones, W.C., 1991. Monthly Variations in the Size of Spicules of the Haplosclerid Sponge, Haliclona rosea (Bowerbank), in: Reitner, J., Keupp, H. (Eds.), Fossil and Recent Sponges. Springer, Berlin, Heidelberg, pp. 404–420. 10.1007/978-3-642-75656-6_33

Jones, W.C., 1987. Seasonal variations in the skeleton and spicule dimensions of Haliclona elegans (Bowerbank) sensu Topsent (1887) from two sites in North Wales, in: Jones, W.C. (Ed.), European Contribution to the Taxonomy of Sponges. Sherkin Island Marine Station Publication, Cork (Ireland), pp. 109–129.

Kaneda, T., 1991. Iso-and anteiso-fatty acids in bacteria: Biosynthesis, function, and taxonomic significance. Microbiol Rev 55, 288–302.

Keesing, J.K., Strzelecki, J., Fromont, J., Thomson, D., 2013. Sponges as important sources of nitrate on an oligotrophic continental shelf. Limnol Oceanogr 58, 1947–1958. 10.4319/lo.2013.58.6.1947

Kelly, J.R., Scheibling, R.E., 2012. Fatty acids as dietary tracers in benthic food webs. Mar Ecol Prog Ser 446, 1–22. 10.3354/meps09559

Kelly, Michelle, Sim-Smith, C., 2023. Kingdom Animalia, phylum Porifera (sponges), in: Kelly, M., Mills, S., Terezow, M., Sim-Smith, C., Nelson, W. (Eds.), The Marine Biota of Aotearoa New Zealand. NIWA Biodiversity Memoir, pp. 83–109.

Kersken, D., Kocot, K., Janussen, D., Schell, T., Pfenninger, M., Martínez Arbizu, P., 2018. First insights into the phylogeny of deep-sea glass sponges (Hexactinellida) from polymetallic nodule fields in the Clarion-Clipperton Fracture Zone (CCFZ), northeastern Pacific. Hydrobiologia 811, 283–293. 10.1007/s10750-017-3498-3

Kumala, L., Larsen, M., Glud, R.N., Canfield, D.E., 2021. Spatial and temporal anoxia in single-osculum Halichondria panicea demosponge explants studied with planar optodes. Mar Biol 168. 10.1007/s00227-021-03980-2

Kutti, T., Bannister, R.J., Fosså, J.H., 2013. Community structure and ecological function of deep-water sponge grounds in the Traenadypet MPA-Northern Norwegian continental shelf. Cont Shelf Res 69, 21–30. 10.1016/j.csr.2013.09.011

Landry, Z., Swa, B.K., Herndl, G.J., Stepanauskas, R., Giovannoni, S.J., 2017. SAR202 genomes from the dark ocean predict pathways for the oxidation of recalcitrant dissolved organic matter. mBio 8. 10.1128/mBio.00413-17

Lawson, M.P., Bergquist, P.R., Cambie, R.C., 1984. Fatty acid composition and the classification of the Porifera. Biochem Syst Ecol 12, 375–393. 10.1016/0305-1978(84)90070-X

Legendre, P., Gallagher, E.D., 2001. Ecologically meaningful transformations for ordination of species data. Oecologia 129, 271–280. 10.1007/s004420100716

Lindsay, D.B., 1975. Fatty acids as energy sources. Proceedings of the Nutrition Society 34, 241–248. 10.1079/pns19750045

Litchfield, C., Greenberg, A.J., Noto, G., Morales, R.W., 1976. Unusually high levels of C24−C30 fatty acids in sponges of the class demospongiae. Lipids 11, 567–570. 10.1007/BF02532903

López-Acosta, M., Leynaert, A., Grall, J., Maldonado, M., 2018. Silicon consumption kinetics by marine sponges: An assessment of their role at the ecosystem level. Limnol Oceanogr 63, 2508–2522. 10.1002/lno.10956

Lüskow, F., Riisgård, H.U., Solovyeva, V., Brewer, J.R., 2019. Seasonal changes in bacteria and phytoplankton biomass control the condition index of the demosponge Halichondria panicea in temperate Danish waters. Mar Ecol Prog Ser 608, 119–132. 10.3354/meps12785

Mackenzie, A.K., Pope, P.B., Pedersen, H.L., Gupta, R., Morrison, M., Willats, W.G.T., Eijsink, V.G.H., 2012. Two SusD-like proteins encoded within a polysaccharide utilization locus of an uncultured ruminant bacteroidetes phylotype bind strongly to cellulose. Appl Environ Microbiol 78, 5935–5937. 10.1128/AEM.01164-12

Maldonado, M., López-Acosta, M., Busch, K., Slaby, B.M., Bayer, K., Beazley, L., Hentschel, U., Kenchington, E., Rapp, H.T., 2021. A Microbial Nitrogen Engine Modulated by Bacteriosyncytia in Hexactinellid Sponges: Ecological Implications for Deep-Sea Communities. Front Mar Sci 8. 10.3389/fmars.2021.638505

Meesters, E., Knijn, R., Willemsen, P., Pennartz, R., Roebers, G., van Soest, R.W.. M., 1991. Sub-rubble communities of Curaçao and Bonaire coral reefs. Coral Reefs 10, 189–197. 10.1007/BF00336773

Moitinho-Silva, L., Steinert, G., Nielsen, S., Hardoim, C.C.P., Wu, Y.C., McCormack, G.P., López-Legentil, S., Marchant, R., Webster, N., Thomas, T., Hentschel, U., 2017. Predicting the HMA-LMA status in marine sponges by machine learning. Front Microbiol 8. 10.3389/fmicb.2017.00752

Morganti, T.M., Slaby, B.M., de Kluijver, A., Busch, K., Hentschel, U., Middelburg, J.J., Grotheer, H., Mollenhauer, G., Dannheim, J., Rapp, H.T., Purser, A., Boetius, A., 2022. Giant sponge grounds of Central Arctic seamounts are associated with extinct seep life. Nat Commun 13, 638. 10.1038/s41467-022-28129-7

Mueller, C.E., Larsson, A.I., Veuger, B., Middelburg, J.J., van Oevelen, D., 2014. Opportunistic feeding on various organic food sources by the cold-water coral Lophelia pertusa. Biogeosciences 11, 123–133. 10.5194/bg-11-123-2014

Müller, W.E.G., Jinhe Li, Schröder, H.C., Li Qiao, Xiaohong Wang, 2007. The unique skeleton of siliceous sponges (Porifera; Hexactinellida and Demospongiae) that evolved first from the Urmetazoa during the Proterozoic: A review. Biogeosciences 4, 219–232. 10.5194/bg-4-219-2007

Muyzer, G., De Waal,’ And, E.C., Uitierlinden2, A.G., 1993. Profiling of Complex Microbial Populations by Denaturing Gradient Gel Electrophoresis Analysis of Polymerase Chain Reaction-Amplified Genes Coding for 16S rRNA, APPLIED AND ENVIRONMENTAL MICROBIOLOGY.

O’Brien, P.A., Tan, S., Frade, P.R., Robbins, S.J., Engelberts, J.P., Bell, S.C., Vanwonterghem, I., Miller, D.J., Webster, N.S., Zhang, G., Bourne, D.G., 2023. Validation of key sponge symbiont pathways using genome-centric metatranscriptomics. Environ Microbiol 25, 3207–3224. 10.1111/1462-2920.16509

Oksanen, J., Blanchet, F.G., Friendly, M., Kindt, R., Legendre, P., McGlinn, D., Minchin, P.R., O’Hara, R.B., Simpson, G.L., Solymos, P., Stevens, H.H., Szoecs, E., Wagner, H., 2017. vegan: Community ecology package.

Pachiadaki, M.G., Sintes, E., Bergauer, K., Brown, J.M., Record, N.R., Swan, B.K., Mathyer, M.E., Hallam, S.J., Lopez-Garcia, P., Takaki, Y., Nunoura, T., Woyke, T., Herndl, G.J., Stepanauskas, R., 2017. Major role of nitrite-oxidizing bacteria in dark ocean carbon fixation, Science.

Perea-Blázquez, A., Davy, S.K., Bell, J.J., 2012. Nutrient utilisation by shallow water temperate sponges in New Zealand. Hydrobiologia 687, 237–250. 10.1007/s10750-011-0798-x

Preston, C.M., Wu, K.Y., Molinski, T.F., DeLong, E.F., 1996. A psychrophilic crenarchaeon inhabits a marine sponge: Cenarchaeum symbiosum gen. nov., sp. nov. Proceedings of the National Academy of Sciences 93, 6241–6246. 10.1073/pnas.93.13.6241

Quast, C., Pruesse, E., Yilmaz, P., Gerken, J., Schweer, T., Yarza, P., Peplies, J., Glöckner, F.O., 2013. The SILVA ribosomal RNA gene database project: Improved data processing and web-based tools. Nucleic Acids Res 41. 10.1093/nar/gks1219

R-Core Team, 2022. R: A language and environment for statistical computing.

Reincke, T., Barthel, D., 1997. Silica uptake kinetics of Halichondria panicea in Kiel Bight. Mar Biol 129, 591–593.

Reiswig, H.M., Kelly, M., 2018. The marine fauna of New Zealand: Euplectellid glass sponges (Hexactinellida, Lyssacinosida, Euplectellidae). NIWA Biodiversity Memoirs 130, 118–123.

Rix, L., de Goeij, J.M., Mueller, C.E., Struck, U., Middelburg, J.J., van Duyl, F.C., Al-Horani, F.A., Wild, C., Naumann, M.S., van Oevelen, D., 2016. Coral mucus fuels the sponge loop in warm- and cold-water coral reef ecosystems. Sci Rep 6, 18715. 10.1038/srep18715

Rossel, S., Peters, J., Charzinski, N., Eichsteller, A., Laakmann, S., Neumann, H., Martínez Arbizu, P., 2024. A universal tool for marine metazoan species identification: towards best practices in proteomic fingerprinting. Sci Rep 14, 1280. 10.1038/s41598-024-51235-z

Ryan, C.G., Clayton, E., Griffin, W.L., Sie, S.H., Cousens, D.R., 1988. SNIP, a statistics-sensitive background treatment for the quantitative analysis of PIXE spectra in geoscience applications. Nucl Instrum Methods Phys Res B 34, 396–402. 10.1016/0168-583X(88)90063-8

Sampath, H., Ntambi, J.M., 2004. Polyunsaturated fatty acid regulation of gene expression. Nutr Rev 62, 333–339. 10.1301/nr.2004.sept.333-339

Savitzky, Abraham., Golay, M.J.E., 1964. Smoothing and Differentiation of Data by Simplified Least Squares Procedures. Anal Chem 36, 1627–1639. 10.1021/ac60214a047

Schreiber, A., Wörheide, G., Thiel, V., 2006. The fatty acids of calcareous sponges (Calcarea, Porifera). Chem Phys Lipids 143, 29–37. 10.1016/j.chemphyslip.2006.06.001

Simion, P., Philippe, H., Baurain, D., Jager, M., Richter, D.J., Di Franco, A., Roure, B., Satoh, N., Quéinnec, É., Ereskovsky, A., Lapébie, P., Corre, E., Delsuc, F., King, N., Wörheide, G., Manuel, M., 2017. A large and consistent phylogenomic dataset supports sponges as the sister group to all other animals. Current Biology 27, 958–967. 10.1016/j.cub.2017.02.031

Soetaert, K., Petzoldt, T., Meysman, F.J.R., 2010. marelac: Tools for Aquatic Sciences. R package version 2.1.

Southwell, M.W., 2007. Sponges impacts on coral reef nitrogen cycling, Key Largo, Florida (PhD Thesis). University of North Carolina, Chapel Hill.

Southwell, M.W., Weisz, J.B., Martens, C.S., Lindquist, N., 2008. *In situ* fluxes of dissolved inorganic nitrogen from the sponge community on Conch Reef, Key Largo, Florida. Limnol Oceanogr 53, 986–996. 10.4319/lo.2008.53.3.0986

Spalding, M.D., Fox, H.E., Allen, G.R., Davidson, N., Ferdaña, Z.A., Finlayson, M., Halpern, B.S., Jorge, M.A., Lombana, A., Lourie, S.A., Martin, K.D., McManus, E., Molnar, J., Recchia, C.A., Robertson, J., 2007. Marine ecoregions of the world: A bioregionalization of coastal and shelf areas. Bioscience 57, 573–583. 10.1641/B570707

Spector, A.A., Yorek, M.A., 1985. Membrane lipid composition and cellular function. J Lipid Res 26, 1015–1035.

Steinert, G., Busch, K., Bayer, K., Kodami, S., Arbizu, P.M., Kelly, M., Mills, S., Erpenbeck, D., Dohrmann, M., Wörheide, G., Hentschel, U., Schupp, P.J., 2020. Compositional and Quantitative Insights Into Bacterial and Archaeal Communities of South Pacific Deep-Sea Sponges (Demospongiae and Hexactinellida). Front Microbiol 11. 10.3389/fmicb.2020.00716

Stratmann, T., Mevenkamp, L., Sweetman, A.K., Vanreusel, A., van Oevelen, D., 2018. Has phytodetritus processing by an abyssal soft-sediment community recovered 26 years after an experimental disturbance? Front Mar Sci 5, 59. 10.3389/fmars.2018.00059

Sun, X., Kop, L.F.M., Lau, M.C.Y., Frank, J., Jayakumar, A., Lücker, S., Ward, B.B., 2019. Uncultured Nitrospina-like species are major nitrite oxidizing bacteria in oxygen minimum zones. ISME Journal 13, 2391–2402. 10.1038/s41396-019-0443-7

Thiel, V., Blumenberg, M., Hefter, J., Pape, T., Pomponi, S., Reed, J., Reitner, J., Wörheide, G., Michaelis, W., 2002. A chemical view of the most ancient metazoa - Biomarker chemotaxonomy of hexactinellid sponges. Naturwissenschaften 89, 60–66. 10.1007/s00114-001-0284-9

Vacelet, J., 1975. Étude en microscopie électronique de l’association entre bactéries et Spongiaires du genre Verongia (Dictyoceratida). J Microsc Biol Cell 23, 271–288.

van Deenen, L.L.M., 1966. Phospholipids and biomembranes. Prog Chem Fats Other Lipids 8, 1–127. 10.1016/0079-6832(66)90003-6

Van Duyl, F.C., Hegeman, J., Hoogstraten, A., Maier, C., 2008. Dissolved carbon fixation by sponge-microbe consortia of deep water coral mounds in the northeastern Atlantic Ocean. Mar Ecol Prog Ser 358, 137–150. 10.3354/meps07370

van Duyl, F.C., Lengger, S.K., Schouten, S., Lundälv, T., van Oevelen, D., Müller, C.E., 2020. Dark CO2 fixation into phospholipid-derived fatty acids by the cold-water coral associated sponge Hymedesmia (Stylopus) coriacea (Tisler Reef, NE Skagerrak). Marine Biology Research 16, 1–17. 10.1080/17451000.2019.1704019

Van Soest, R.W.M., Boury-Esnault, N., Vacelet, J., Dohrmann, M., Erpenbeck, D., De Voogd, N.J., Santodomingo, N., Vanhoorne, B., Kelly, M., Hooper, J.N.A., 2012. Global diversity of sponges (Porifera). PLoS One 7, e35105. 10.1371/journal.pone.0035105

Webster, N.S., Taylor, M.W., 2012. Marine sponges and their microbial symbionts: Love and other relationships. Environ Microbiol 14, 335–346. 10.1111/j.1462-2920.2011.02460.x

Weiss, R.F., 1970. The solubility of nitrogen, oxygen and argon in water and seawater. Deep-Sea Research and Oceanographic Abstracts 17, 721–735. 10.1016/0011-7471(70)90037-9

Zea, S., 1993. Cover of sponges and other sessile organisms in rocky and coral reef habitats of Santa Marta, Colombian Caribbean Sea. Caribb J Sci 29, 75–88.

